# Single-cell multidimensional profiling of tumor cell heterogeneity in supratentorial ependymomas

**DOI:** 10.1101/2024.08.07.607066

**Authors:** Daeun Jeong, Sara G. Danielli, Kendra K. Maaß, David R. Ghasemi, Svenja K. Tetzlaff, Ekin Reyhan, Carlos Alberto Oliveira de Biagi-Junior, Sina Neyazi, Andrezza Nascimento, Rebecca Haase, Costanza Lo Cascio, Bernhard Englinger, Li Jiang, Cuong M. Nguyen, Alicia-Christina Baumgartner, Sophia Castellani, Jacob S. Rozowsky, Olivia A. Hack, McKenzie L. Shaw, Daniela Lotsch-Gojo, Katharina Bruckner, Stefan M. Pfister, Marcel Kool, Tomasz J. Nowakowski, Johannes Gojo, Lissa Baird, Sanda Alexandrescu, Kristian W. Pajtler, Varun Venkataramani, Mariella G. Filbin

## Abstract

Supratentorial ependymomas are aggressive childhood brain cancers that retain features of neurodevelopmental cell types and segregate into molecularly and clinically distinct subgroups, suggesting different developmental roots. The developmental signatures as well as microenvironmental factors underlying aberrant cellular transformation and behavior across each supratentorial ependymoma subgroup are unknown. Here we integrated single cell- and spatial transcriptomics, as well as *in vitro* and *in vivo* live-cell imaging to define supratentorial ependymoma cell states, spatial organization, and dynamic behavior within the neural microenvironment. We find that individual tumor subgroups harbor two distinct progenitor-like cell states reminiscent of early human brain development and diverge in the extent of neuronal or ependymal differentiation. We further uncover several modes of spatial organization of these tumors, including a high order architecture influenced by mesenchymal and hypoxia signatures. Finally, we identify an unappreciated role for brain-resident cells in shifting supratentorial ependymoma cellular heterogeneity towards neuronal-like cells that co-opt immature neuronal morphology and invasion mechanisms. Collectively, these findings provide a multidimensional framework to integrate transcriptional and phenotypic characterization of tumor heterogeneity in supratentorial ependymoma and its potential clinical implications.

## Main

Brain cancers are the leading cause of cancer-related deaths in children and adolescents, surpassing leukemia^1^. Deciphering the developmental correlates is pivotal to therapeutic advancement as malignant cells retain molecular and phenotypic properties of their progenitor origins^2–4^. Supratentorial ependymomas (ST-EPNs) are childhood brain tumors that occur in the central nervous system, with varied outcomes and limited available therapies^5–8^. While radial glia cells during early brain development have been proposed as the cells of origin for ST-EPNs^9–11^, differences across species underscore the challenge in extrapolating findings from these studies in mice. Moreover, recent genome-wide DNA methylation profiling have classified ST-EPNs into multiple subgroups with distinct fusion genes and patient outcomes^12,13^. These include the canonical ST-ZFTA subgroup characterized by the fusion of NFkB pathway regulator *RELA* with the zinc finger translocation associated *ZFTA* (*ZFTA-RELA*), and the non-canonical ST-ZFTA subgroups (ZFTA-Clusters 1 to 4), which harbor *ZFTA-RELA* fusions or fusions between *ZFTA* and other partner genes. Additionally, the ST-YAP1 subgroup is enriched for fusions of Hippo effector *YAP1*. It remains unknown whether these subgroups have distinct cellular origins and composition of malignant cell states leading to these differences in outcomes and varying resistance to therapy. To address these questions, previous studies have leveraged single cell and single nucleus RNA-sequencing (sc/snRNA-seq)^14,15^ to characterize the molecular cell states of ST-EPN tumors based on distinct gene expression profiles. However, these studies were limited by patient number and did not include the non-canonical ST-ZFTA subgroups, which led to many cell states being sample- or subgroup-specific. These limitations highlight the need for a comprehensive study examining tumors across all subgroups of ST-EPNs.

In addition to the cell-intrinsic properties, it has recently become clear how cancer cell diversity is further influenced by their microenvironment, where interactions with adjacent cells and non-genetic factors such as deprivation of oxygen or nutrients augment cellular variability and malignant phenotypes^16^. For instance, several studies have shown that the interactions between tumor cells and neurons in gliomas promote proliferation and invasion^17–19^, while regional hypoxia drives transcriptional adaptation and genomic instability in tumor cells^20,21^. Recent advances in spatial transcriptomics have overcome the loss of spatial information in scRNA-seq analyses, thereby revealing the intricate ecosystem within various tumors^22–24^. In addition, as these two methods still require cells to be fixed in time and space, live-cell imaging of gliomas has proven effective for capturing the dynamic nature of tumor cells within their microenvironment while preserving cell viability, morphology, and function^25^. Such multilayered analysis linking cell state to microenvironmental influences, dynamic morphology and behavior of malignant cells have yet to be done in ST-EPN tumors.

Here, we integrate sc/snRNAseq, spatial transcriptomics, and *in vitro* and *in vivo* live-cell imaging to provide a multidimensional framework to characterize tumor heterogeneity in ST-EPNs. We first delineate the developmental correlates of different ST-EPN subgroups and find that malignant cells reflect two distinct temporally restricted progenitors from early human embryonic cortical development and exhibit varying degrees of neuronal or ependymal differentiation. Furthermore, we uncover global spatial organization patterns driven by hypoxia, as well as local tumor niches that reveal cell state-driven localization. Lastly, we find that cell states display distinct morphological and migratory patterns, with neuronal-like cancer cells being the most invasive tumor compartment by adopting mechanisms of neuronal migration during development. We demonstrate the crucial role of the neural microenvironment in promoting plasticity of cells toward this neuronal lineage. Taken together, we unveil the extensive tumor heterogeneity in ST-EPNs by shedding light on their developmental cell states, patterns of spatial localization, and cellular morphology and behavior, thereby opening new potential avenues for therapeutic interventions.

## Results

### Supratentorial ependymoma (ST-EPN) subgroups are transcriptionally distinct

To comprehensively characterize the cellular tumor heterogeneity of supratentorial ependymomas (ST-EPN), we compiled a patient cohort of 44 patient tumors encompassing ZFTA-RELA (*n* = 20), ZFTA-Clusters 1-4 (*n* = 20) and ST-YAP1 (*n* = 4) subgroups. ZFTA-RELA, ZFTA-Cluster 1 and ZFTA-Cluster 3 harbored canonical *ZFTA-RELA* fusions, ZFTA-Cluster 2 and ZFTA-Cluster 4 harbored non-canonical *ZFTA* fusion types, and ST-YAP1 harbored fusions of *YAP1* (**Figure 1a-b, Table S1)**. Orthotopic patient-derived xenografts (ZFTA-RELA PDX *n*=3) and patient-derived cell lines (ZFTA-RELA *n* = 3) were also included. We performed deep full-length single-cell transcriptomics on frozen (single-nucleus RNA-sequencing, snRNA-seq) and fresh (single-cell RNA-sequencing, scRNA-seq) tumor specimens, and retained a total of 7,429 high-quality cells, which include a mix of malignant tumor cells and non-malignant cell types derived from the tumor microenvironment. To distinguish malignant cells from non-malignant cell types, we used three complementary approaches: genome-wide copy-number alterations, similarity between the transcriptome of each single-cell and a reference collection of healthy cell types, and expression of canonical marker genes (**Extended Data Figure 1a-b**). We retained 6,933 high-quality malignant cells (mean of 5,511 genes per cell/nucleus; **Extended Data Figure 1c, Table S1**), which we processed uniformly and co-embedded into a shared low-dimensional space.

**Figure 1.**
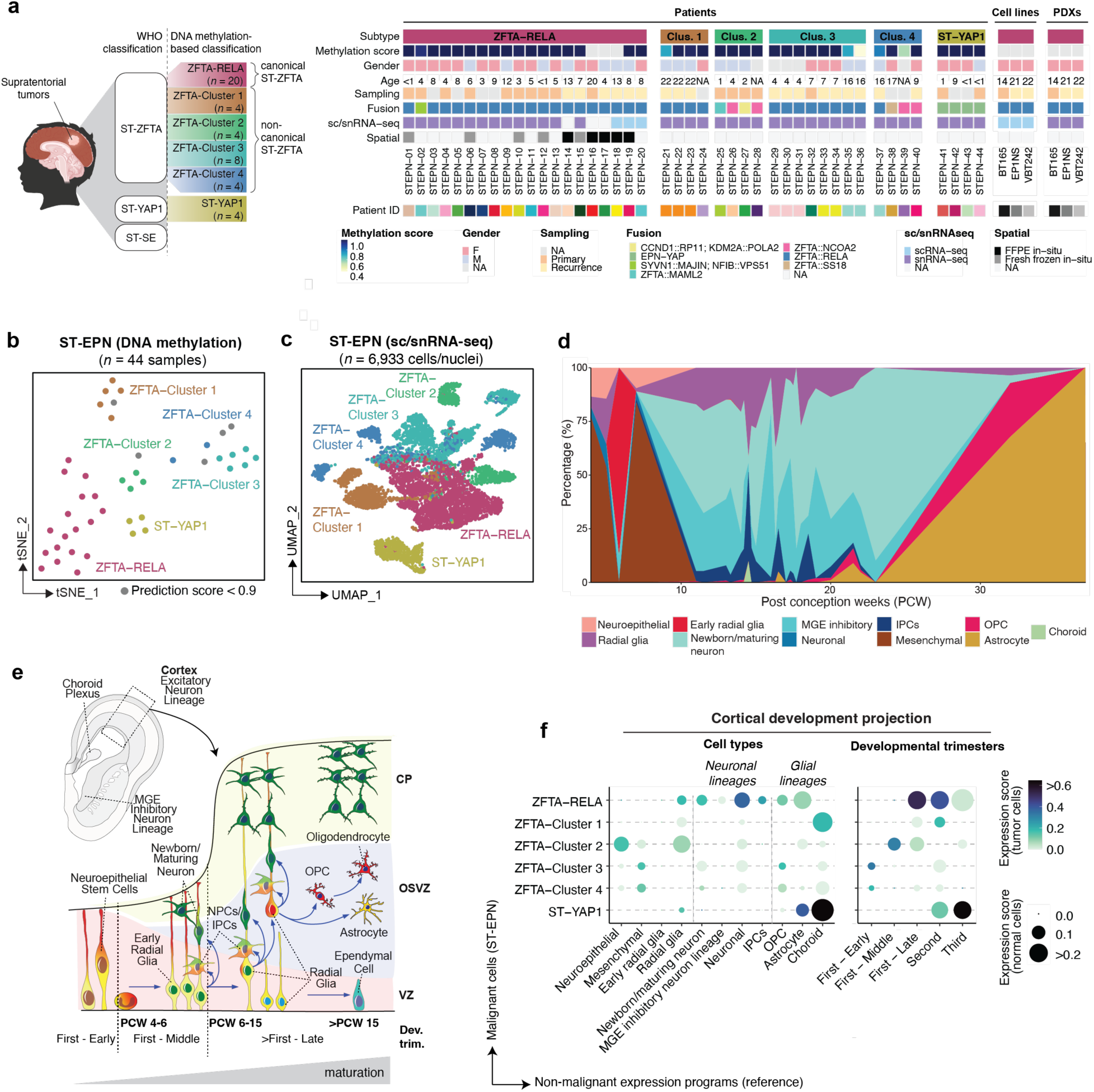
Molecular signatures of ST-EPN subgroups differentially project to human cortical development. **(a)** Oncoplot depicting ST-EPN samples profiled in this study. **(b)** tSNE plot of ST-EPN samples profiled by DNA methylation profiles and colored by molecular subgroup. Samples with a methylation prediction score < 0.9 are indicated in grey. (**c**) UMAP plot of malignant ST-EPN cells profiled by sc/snRNA-seq and colored by molecular subgroup. **(d)** Percentage of cell types (y-axis) present during the first, second and third trimester of human cortical development (x-axis). Data integrated and reanalyzed from two published scRNA-seq datasets of the human developing cortex^29,30^ **e)** Simplified overview of cell types present during human cortical development. **(f)** Projections of ST-EPN subgroups (y-axis) onto cell types or developmental time points found across the developing human cortex (x-axis). First-Early: post-conception week (PCW) 4, First-Middle: PCW 5-6, First-Late: PCW 7-12. Color scale depicts expression score of normal cell type gene expression in tumor cells, and dot size represents expression score of tumor cell type gene expression in normal cells. PCW, post-conceptional week; dev. trim, developmental trimesters; NPCs, neural-progenitor cells; IPCs, intermediate progenitor cells; OPC, oligodendrocyte progenitor cells; MGE, medial ganglionic eminence.

As DNA methylation profiles utilized to categorize these tumors are considered epigenetic fingerprints of cellular origins in both development and tumorigenesis^26,27^, we hypothesized that ST-EPN tumors would correspondingly be transcriptionally distinct. Accordingly, we find that the molecular subgroup classification drives clustering of all malignant cells (**Figure 1b-c**), consistent with previous observations^28^. We next examined whether the inter-tumoral differences observed across subgroups reflect different developmental origins and compared transcriptomic profiles of malignant cells with normal tissues transcriptomes. We first assembled a single-cell reference atlas of the developing human cerebral cortex from two published scRNA-seq datasets (**Figure 1d, Extended Data Figure 1d**)^29,30^ that includes all three prenatal trimesters of embryonic and fetal development from post conception weeks (PCWs) 4 to 37 (**Table S2-S3**). Next, we aggregated the gene expression profiles of individual ST-EPN cells by molecular subgroup, resulting in a pseudobulked gene expression matrix which we projected onto the generated reference atlas^29,30^. ST-EPN tumor subgroups showed significant differences in their correlations to immature cell types of the developing human cerebral cortex (**Figure 1e-f)**. While canonical ZFTA-RELA tumors most closely resembled cells of the neuronal lineage, ZFTA-Cluster 2 tumors preferentially mapped to early progenitors within the brain, including neuroepithelial stem cells and radial glia cells. In contrast, ZFTA-Cluster 3 and 4 tumors mapped to mesenchymal cells found in the early first trimester, and ZFTA-Cluster 1 and ST-YAP1 tumors projected mostly onto cells originally classified as choroid (**Figure 1f**). Because the choroid plexus in the reference dataset consists mostly of gene programs related to cilia processes, and ST-EPN tumors displayed strong *FOXJ1,* a typical ependymal marker^31^, and no *TTR* expression, a typical marker for choroid plexus tissue^32^ (**Extended Data Figure 1e**), we considered ependymal and choroid cells interchangeably in this context. Interestingly, ZFTA-Cluster 2-4 tumors generally mapped to early first trimester prenatal cell types, while ZFTA-Cluster 1, canonical ZFTA-RELA and ST-YAP1 tumors mapped to later embryonic timepoints (**Figure 1f**).

In summary, our analysis reveals that molecularly distinct ST-EPN subgroups align with temporally restricted cells across human cortical development. Moreover, they are defined by the specificity and extend of their maturation along either a neuronal or glial lineage trajectory.

### The cellular hierarchy of ST-EPN tumors mirrors human cortical development

To identify specific cell states across ST-EPN tumor subgroups, we next moved from pseudobulk projections to comparing single cell expression identified in each sample by non-negative matrix factorization (NMF)^33^. This analysis revealed eight clusters of recurrent programs across ST-EPN tumors (**Figure 2a**), termed metaprograms, which reflect relative cellular states. We annotated each metaprogram based on the expression of known cell type-specific marker genes (**Table S4**), projection onto developing human cerebral cortex reference atlases (**Extended Data Figure 2a**), and functional analysis using Gene Ontology (GO) enrichment analysis (**Extended Data Figure 2b, Table S5**).

**Figure 2.**
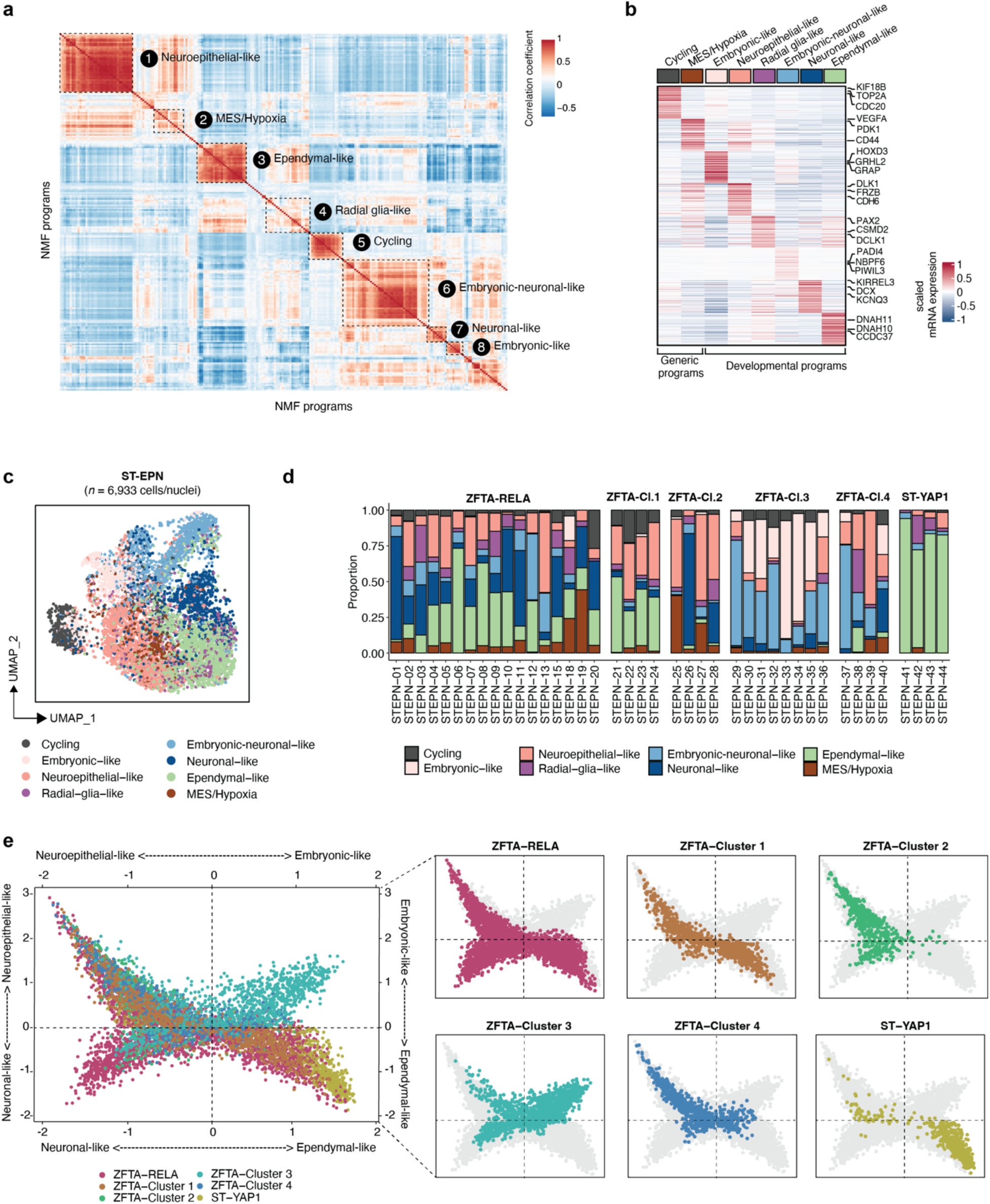
Intra-tumoral heterogeneity of ST-EPN tumors. **(a)** Heatmap showing Pearson correlation coefficient between individual NMF programs identified within each ST-EPN sample profiled by snRNA-seq. Programs are grouped into eight recurrent metaprograms (dashed lines). **(b)** Heatmap showing relative expression (z-score) of the top 200 genes for each metaprogram across all ST-EPN cells profiled in this study. Selected genes are highlighted. **(c)** UMAP plot of malignant ST-EPN cells colored by identified cellular state. Samples were integrated using harmony^68^. **(d)** Proportion of cellular states in each ST-EPN sample (x-axis). **(e)** Cells state plot of malignant ST-EPN cells. Cells were scored for the metaprograms identified by NMF analysis (top 30 genes) and colored by molecular subgroup.

Two metaprograms were associated with generic cancer cell processes, including a cycling metaprogram (*MKI67, TOP2A*), which is enriched in genes associated with cell proliferation, and a mesenchymal (MES)/Hypoxia (*CD44, VEGFA*) metaprogram, which highly expresses hypoxia-related genes^34^ (**Figure 2b, Table S4**). The remaining six metaprograms were distinctly related to developmental programs, including: (1) embryonic-like (*HOXD3*) enriched in GO terms related to embryonic morphogenesis and organ development but not matching to a particular cell type found in the developing human brain^35^; (2) neuroepithelial-like (*PTGDS*, *TAGLN2)* that strongly project to neuroepithelial stem cells of the developing human cortex; (3) radial glia-like that expresses radial glia genes (*TNC*, *LRIG1*)^36,37^; (4) embryonic-neuronal-like (*PADI4, NBPF6, PIWIL3*) enriched with immature neuronal genes^32^; (5) neuronal-like (*DCX, TUBB3*) enriched in neuronal differentiation-related genes^38,39^; (6) ependymal-like (*DNAH9, DNAH6*) enriched with cilia-related genes^32^(**Figure 2b, Extended Data Figure 2b, Table S4-5**). These metaprograms partially correlated with those identified in posterior fossa, spinal and supratentorial EPNs of previous studies^14^ (**Extended Data Figure 2c**) and corroborated our previous pseudobulk projection onto the developing human cerebral cortex (**Figure 1f, Extended Data Figure 2a**). Metaprogram gene signatures were queried through the Drug Gene Interaction database (DGIdb)^40^ uncovering potential therapeutic targets of relevance, especially in the context of metaprograms correlated with lower survival (**Extended Data Figure 3a-b, Table S6**).

We next examined how the proportion of the identified metaprograms differs across ST-EPN subgroup. While ZFTA-RELA tumors were composed predominantly of neuronal-like and ependymal-like cells, neuronal-like cells were absent in ZFTA-Cluster 1 and ST-YAP1 tumors (**Figure 2c-d**). Conversely, ZFTA-Cluster 2 and ZFTA-Cluster 4 tumors harbored lower frequency of ependymal-like cells. In addition, ZFTA-Cluster 3 tumors displayed a distinct intra-tumoral composition, being mostly composed of embryonic-like and embryonic-neuronal-like subpopulations (**Figure 2d**). This is consistent with the undifferentiated histology of ZFTA-Cluster 3 tumors, previously often diagnosed as sarcoma, CNS embryonal or other primitive tumors^12^.

To summarize the distribution of metaprograms in each ST-EPN subgroup, we scored all single cells for the two most immature (neuroepithelial-like and embryonic-like) and the two more mature (neuronal-like and ependymal-like) lineage-related expression programs and visualized the results as a cell-state plot (**Figure 2e**). We found that the two immature cell states existed in mutually exclusive manner, indicating that when neuroepithelial-like cells were present in ZFTA-RELA, ZFTA-Cluster 1,2 and 4, there were no embryonic-like cells present. Conversely, in ZFTA-Cluster 3 where embryonic-like cells were highly present, neuroepithelial-like cells were much less abundant. This likely indicates two distinct developmental signatures of ST-EPN, with ZFTA-Cluster 3 as a distinct entity from other subgroups. Interestingly, both ZFTA-RELA and ZFTA-Cluster 3 harbor *ZFTA-RELA* fusions (**Figure 1a**). Differential gene expression analysis of subgroups **(Extended Data Figure 4a)** uncovered early development and morphogenic-related gene signatures enriched in ZFTA-Cluster 3, correlating with the higher proportion of embryonic-like cells in these tumors. Moreover, tumors from different subgroups were distinct in the lineages in which they represent. While there was a continuum of gene expression to both neuronal and ependymal differentiation in ZFTA-RELA tumors, the other subgroups only harbored differentiation to one of the mature cell states (**Figure 2e**). In addition, ST-YAP1 tumors only exhibited ependymal differentiation, with very few progenitor cells present. We validated these findings using an external patient cohort^13^ and confirmed that ST-YAP1 patients map onto ependymal signatures, whereas ZFTA-RELA patients onto neuronal-like signatures (**Extended Data Figure 4b-c**).

Overall, our findings suggest that ST-EPN tumors from different molecular subgroups have distinct developmental signatures, with variability in differentiation of neuronal and ependymal lineages. We highlight ZFTA-Cluster 3 especially as a distinct entity in progenitor and lineage transcriptional profiles.

### Spatial mapping of ZFTA-RELA tumor cell states

We next investigated how these identified tumor cell states are spatially organized in the tissue, and whether recurrent patterns of localization exist across tumors. To do so, we performed 10X Genomics Xenium *in situ* transcriptomics on a total of 23 tumor sections from 10 ZFTA-RELA patients to allow for subcellular mapping of 354 target genes (**Table S7**). We focused on tumors of the canonical ZFTA-RELA subgroup as they encompass the majority of ST-EPNs^13^. We used a combination of custom-based gene panel curated from our scRNA-seq dataset containing marker genes of identified tumor subpopulations and fusion-specific probes to detect fusion transcripts, and a human brain gene panel containing normal cell type-specific genes (**Figure 3a**).

**Figure 3.**
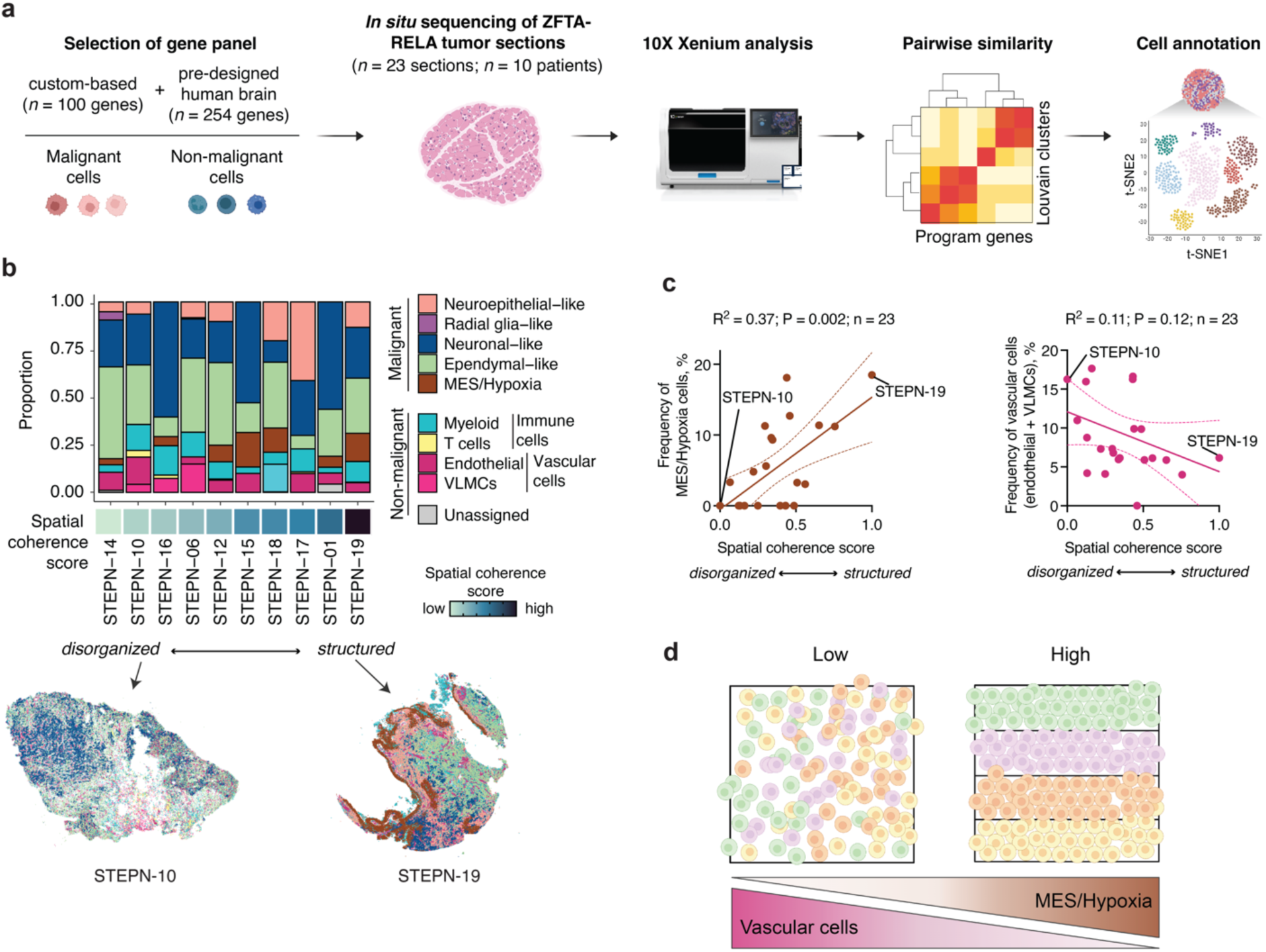
Global spatial architecture of ZFTA-RELA tumors. **(a)** Schematic of 10X Xenium *in situ* workflow. **(b)** Proportion of tumor metaprograms and non-malignant cell types (y-axis) in the ZFTA-RELA patient tumors profiled by 10X Xenium (x-axis). For patients with multiple tumor sections analyzed, the average proportion is shown. Samples are ordered by increasing spatial coherence score. Below are spatial maps of two sections, demonstrating differences in spatial organization between disorganized (STEPN-10) and structured (STEPN-19) tumors. **(c)** Correlation between the spatial coherence score and the percentage of MES/Hypoxia-like or vascular cells in ZFTA-RELA tumor sections profiled by 10X Xenium. Data points are interpolated with a simple linear regression. Dashed line represents the 95% confidence intervals. The goodness of fit (R^2^), P-value and number of data points are displayed at the top. Dots representing the tumor sections depicted in Fig. 3b are labeled. **(d)** Graphical summary of the global spatial organization of ZFTA-RELA tumors.

We first annotated each cell *in situ* by assessing similarities between the top markers of the spatially identified Louvain clusters and previously defined tumor cell states and/or non-malignant cell type marker genes. This analysis identified on average 152,541 cells per tumor section to identify nine spatial cell types/states, including four non-malignant cell types and five malignant cell states (**Figure 3b, Extended Data Figure 5a, 6a-b**). Importantly, by designing fusion-specific probes for each patient, we observed that malignant cells highly expressed *ZFTA-RELA* fusions (**Extended Data** Figure 6c). For malignant cells, we found consistent presence of the neuroepithelial-like, ependymal-like, neuronal-like and MES/Hypoxia states identified above by sc/snRNA-seq, but not of the radial glia-like state, which was detected only in one tumor section derived from one patient (**Figure 3b**, **Extended** Data Figure 6b). Similarly, oligodendrocytes were identified only in a single tumor section derived from a unique patient and were therefore excluded from downstream analyses. Overall, the proportions of tumor cell states within samples were highly correlated to proportions detected by sc/snRNA-seq (R^2^ = 0.901; *P* = 9.12e-05, Extended Data Figure 6a), and was highly consistent across consecutive sections from individual patients, confirming high reproducibility of this technology (**Extended Data Figure 6b**).

### ZFTA-RELA tumors segregate into globally structured and disorganized patterns

After annotating spatially distributed cell states, we observed that tumors segregated into two global organization patterns – (1) high compartmentalization of cell states, which we term *structured* tumors, and (2) cell states/types scattered throughout the section, which we term *disorganized* tumors. (**Extended Data Figure 5a**). We quantified the degree of spatial organization of each tumor section by measuring the degree by which cells from the same state/type are surrounded by other cells of the same state/type, termed *spatial coherence score* (**Extended Data Figure 6d**). Sections that appear *structured* will therefore display high spatial coherence scores; inversely, sections that appear *disorganized* will display low spatial coherence scores. By applying our algorithm to 23 tumor sections, we successfully distinguished between tumor sections that were *structured* (e.g. STEPN-19), and those that were *disorganized* (e.g. STEPN-10; **Figure 3b**).

To determine whether presence of a particular tumor state/type correlates with spatial organization, we performed linear regression analyses between the spatial coherence score of each section and the frequency of cells observed for each state/type. This analysis revealed that higher spatial coherence scores were positively correlated with the frequency of MES/Hypoxia cells (*R*^2^ = 0.37, *P* = 0.002, *n* = 23, **Figure 3c, Table S8**), suggesting that the presence of this malignant cell state is associated with *structured* tumors. We reasoned that vascular perfusion might decrease hypoxia, and therefore revert this structural organization. Indeed, ZFTA-RELA tumors that were more vascularized contained less MES/Hypoxia cells (*R*^2^ = 0.27, *P* = 0.011, *n* = 23), and were generally more disorganized (*R*^2^ = 0.11, *P* = 0.12, *n* = 23, **Figure 3c**, **Extended Data Figure 6e**). These findings indicate that tumor areas that are well perfused display less hypoxia and tend to be spatially *disorganized*, whereas tumors that contain hypoxic regions tend to be more *structured* (**Figure 3d**). In addition, while spatial coherence was highly variable across different samples, relapsed tumors in general were more *structured*, in line with higher frequency of MES/Hypoxia cells (**Extended Data Figure 6f-g**). This indicates that higher spatial organization of ZFTA-RELA tumors correlates with more aggressive tumors.

Overall, our results suggest that ZFTA-RELA tumors are highly compartmentalized tumors driven by the presence of the MES/hypoxia cellular states, and that tumor recurrence is correlated with higher spatial organization.

### ZFTA-RELA tumors are locally organized into recurrent tumor niches

We next sought to understand the local spatial relationships between cell states and/or cell types by identifying groups of cells with similar neighborhoods within the tumor, or *spatial niches*. This analysis, which identifies cells that have similar local neighborhoods within a tissue, showed demarcated regions that were both transcriptionally and morphologically distinct (**Figure 4a-b**, **Extended Data Figure 7a**). We identified niches containing ependymal rosettes composed of ependymal cells, and perivascular rosettes composed of endothelial cells (**Figure 4a)**, demonstrating the direct correlation between transcriptional cellular states and morphology. In addition, regions with clear cytoplasm adjacent to dense nuclei regions overlayed directly with neuronal-like and ependymal-like cells, respectively, present in distinct spatial niches (**Extended Data Figure 7a**).

**Figure 4.**
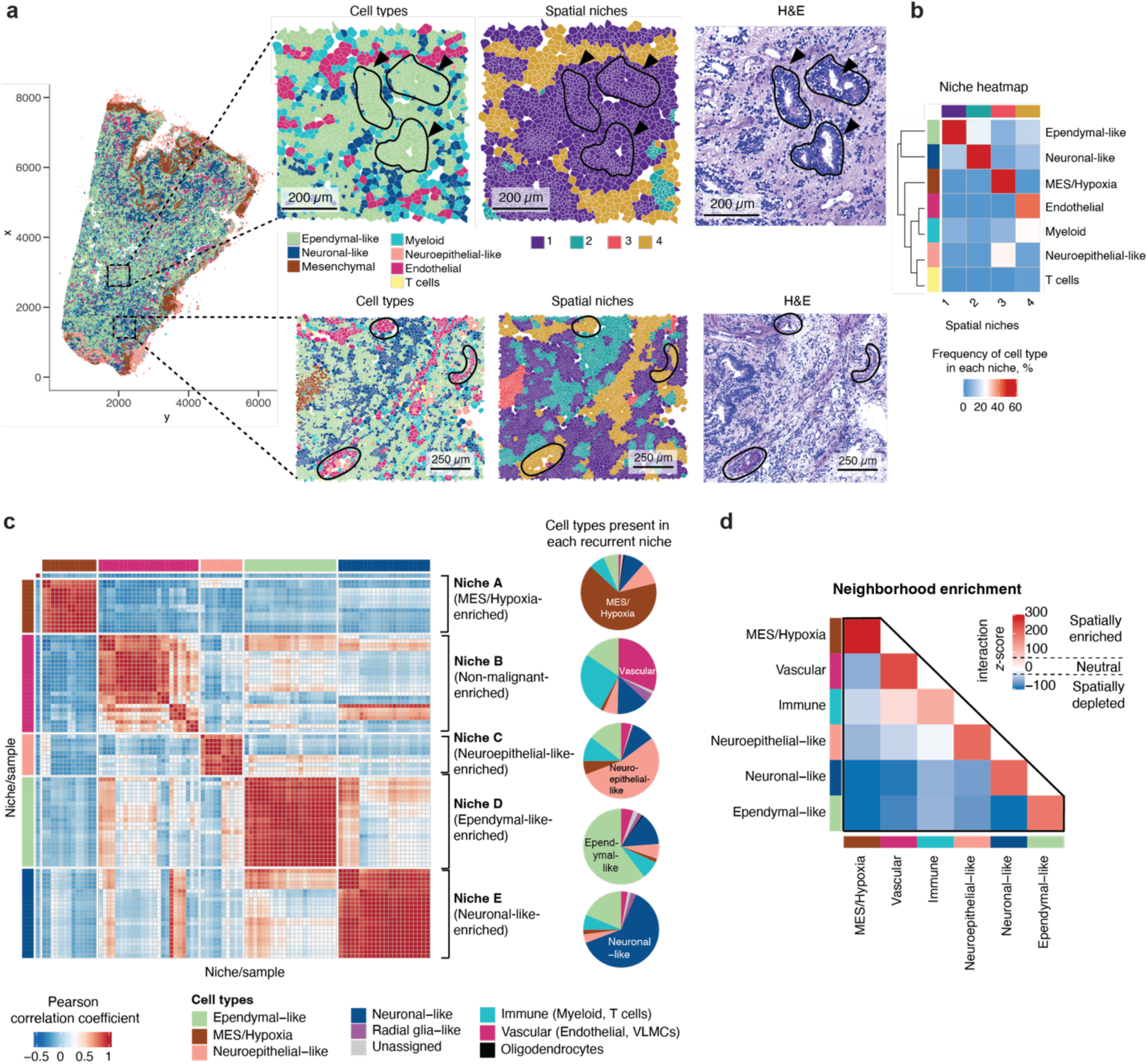
Local spatial architecture of ZFTA-RELA tumors and cellular interactions. **(a)** Overlay of 10X Xenium metaprogram assignment, spatial niches, and H&E staining for ZFTA-RELA tumor section (STEPN-17), showing morphological differences across regions and spatial niches corresponding to different tumor metaprograms. **(b)** Heatmap of corresponding section (STEPN-17), depicting correlation between spatial niche (x-axis) and tumor metaprogram (y-axis), where color scale represents frequency of each cell state/type in each niche. **(c)** Recurrent niche programs identified across the 23 ZFTA-RELA tumor sections analyzed by 10X Xenium. Niches are annotated based on the most frequent cell types (displayed on the right side in the pie charts). **(d)** Neighborhood enrichment analysis showing the tendency of the populations to sit in spatial proximity. The mean interaction z-scores calculated across individual tumor sections (*n* = 23) is shown.

Next, we aimed at uncovering spatial niches that were consistently represented across multiple tumor sections, which we termed *recurrent niches* (**Extended Data Figure 7b**). We overall identified five recurrent niches (A-E): (A) MES/Hypoxia-enriched; (B) non-malignant-enriched; (C) neuroepithelial-like-enriched; (D) ependymal-like-enriched; and (E) neuronal-like-enriched **(Figure 4c**). Most recurrent niches were predominantly composed of one malignant cell state (**Figure 4c**), which indicates that malignant cell states preferentially co-localize with themselves. This was observed in both structured and disorganized tumors (**Extended Data Figure 7c**). We also performed an orthogonal method to determine neighborhood enrichment within each tumor section. This analysis corroborated findings from analysis of recurrent niches, where most malignant cell states preferentially clustered with themselves (**Figure 4d**). This was especially evident for MES/Hypoxia cells which exhibited the strongest neighborhood interactions, (**Figure 4d).** In addition, non-malignant immune (myeloid and T cells) and vascular cells showed preferential co-localization (**Figure 4d**), as observed in recurrent niche B, enriched in both vascular and immune cell types (**Figure 4c**). Overall, we uncovered local spatial patterns of cell states across ZFTA-RELA tumors, with a common observation of malignant cell states preferentially co-localizing with themselves.

### Superimposing molecular and morphological characteristics with function in ZFTA-RELA tumors

Recent studies have revealed that neuronal activity can promote the proliferation and invasion of distinct molecularly defined tumor subpopulations through paracrine and synaptic signaling^18,19,41^. Having observed that malignant cell states with distinct gene expression profiles in ZFTA-RELA tumors overlay with morphologically distinct tumor regions by H&E staining, we next examined whether cell states in ZFTA-RELA tumors also have distinct morphological and functional behavior influenced by neighboring neurons and glial cells.

We incorporated two complimentary models using patient-derived human ZFTA-RELA cells: *in vitro* co-culture of tumor cells with primary rat neurons and astrocytes, and *in vivo* patient-derived xenograft (PDX) models. We first assessed morphological features of tumor cells grown under both conditions. In both models, we observed tumor cells exhibiting immature neuron-like membrane protrusions resembling tumor microtubes (TMs)^42^ classified into four distinct morphological subtypes (**Figure 5a**): cells without processes (0 TM), cells with 1-3 short primary processes, cells with more than three short primary processes and cells with 1-3 long primary processes (longer than 100 μm). We next aimed at overlaying these morphologies with the molecular ZFTA-RELA cell state classification and selected markers from our sc/snRNA-seq atlas (**Table S4**) as well as known neurodevelopmental markers. Subsequently, we performed immunofluorescence staining together with high-resolution light microscopy to confirm the specificity of our approach. We identified: Vimentin+/Nestin+ neuroepithelial-like cells, classified as cells with either no or 1-3 short TMs; S100B+/GFAP+ EPN-glial-like cells, classified as cells with >3 short TMs; and beta-III-tubulin+/DCX+ neuronal-like cells, classified as cells with 1-3 long TMs (**Figure 5b, Extended Data Figure 8a**). In addition, we confirmed the validity of this integrated morpho-molecular classification system by 10X Xenium on ZFTA-RELA cells in co-culture and correlating gene expression derived from individual cells of representative morphologies with gene signatures of tumor metaprograms (**Extended Data Figure 8b-d**). Collectively, we were able to overlay molecularly defined ZFTA-RELA cellular states with distinct morphologies in both *in vivo* and *in vitro* coculture models.

**Figure 5.**
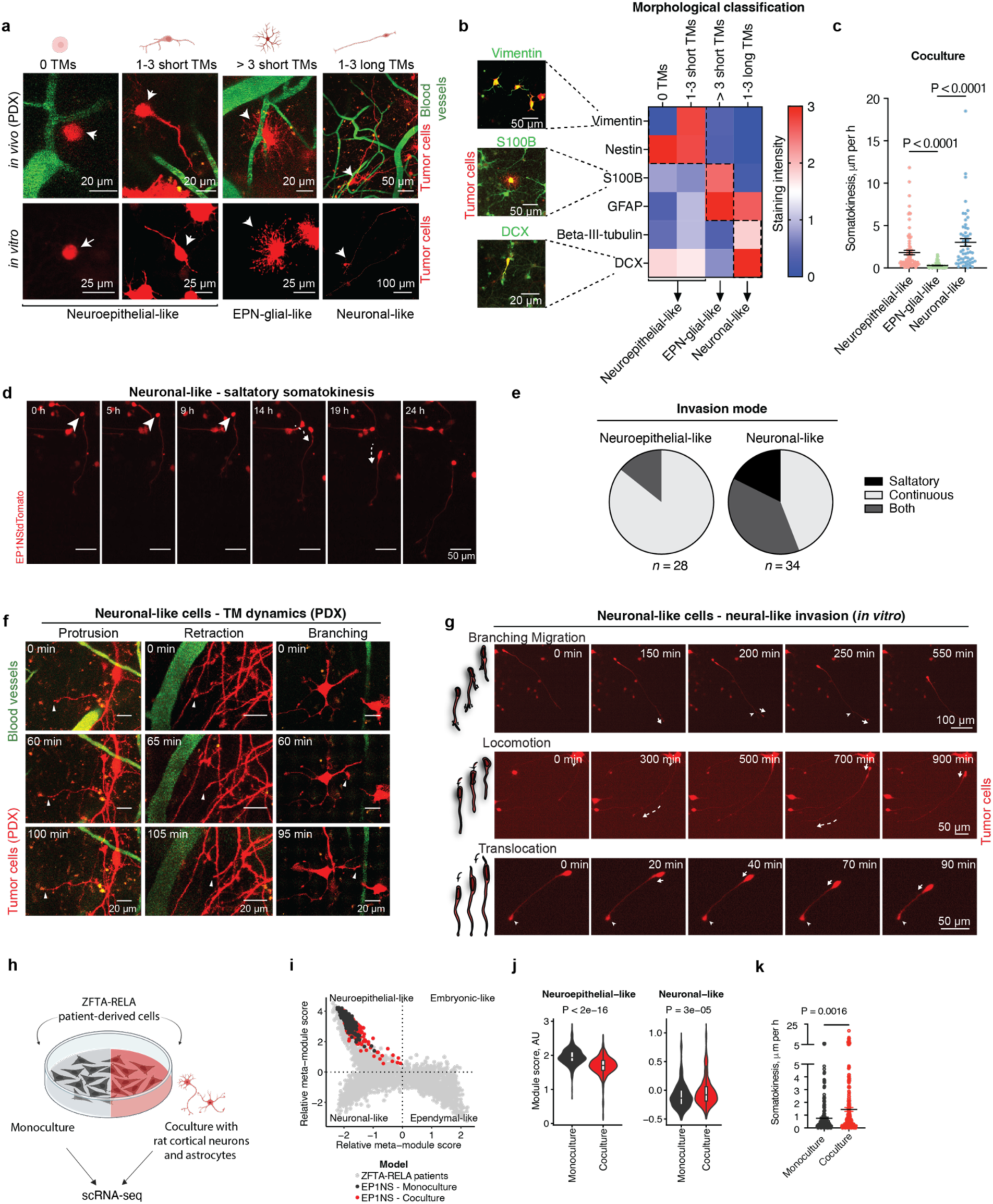
Morpho-molecular classification of ZFTA-RELA cell populations. **(a)** Morphological characterization of ZFTA-RELA cells (EP1NS) in an *in vivo* PDX (top row) and *in vitro* coculture model (bottom row). ZFTA-RELA cells express endogenous tdTomato. *In vivo* PDX images were acquired using in-vivo two-photon microscopy; blood vessels are shown in green. ZFTA-RELA cells are classified as: round cells without TMs (neuroepithelial-like, left), cells with 1-3 short TMs (neuroepithelial-like, middle left), cells with more than 3 short TMs (EPN-glial-like, middle right) and cells with 1-3 long TMs (neuronal-like, right). **(b)** Representative immunofluorescence images and mean intensity of signal of an *in vitro* co-culture model of ZFTA-RELA cells expressing tdTomato, categorized by morphological cell subtypes. Data represent the mean intensity value of *n* = 234 cells acquired from *n* = 3 independent biological replicates per marker. **(c)** Somatokinesis of an *in vitro* coculture model of ZFTA-RELA cells in µm per hour, determined using timelapse imaging in-vitro. Cells are categorized by morphological subtype (*n* = 235 cells acquired from *n* = 3 independent biological replicates). Individual data points with mean±SE are shown. *P* values calculated by Kruskal-Wallis test. **(d)** *In vivo* time-lapse imaging of tumor microtube dynamics in a PDX model of ZFTA-RELA cells expressing tdTomato. Arrowheads point to the ends of tumor microtubes. Blood vessels are shown in green. **(e)** Quantification of invasion modes across ZFTA-RELA cells classified as neuroepithelial-like or neuronal-like based on morphology. The total number of cells is displayed at the bottom. **(f)** In-vivo time-lapse imaging of tumor microtube dynamics in an *in vivo* PDX model of ZFTA-RELA cells expressing tdTomato. Arrowheads point to the ends of tumor microtubes. Blood vessels are shown in green. Different forms of TM dynamics, such as protrusion, retraction, branching and TM genesis using live cell timelapse imaging *in vitro*. Arrows indicate direction of TM movement. Arrowhead pointing to soma, out of which a new primary TM grows. **(g)** Neuronal-like invasion patterns of an *in vitro* model of ZFTA-RELA cells expressing tdTomato shown in a schematic and live-cell timelapse imaging examples. Arrows indicate direction of TM movement. Arrowhead pointing to branching point (branching migration), cell soma (locomotion) or stable TM tip (translocation). **(h)** Experimental design to test the compositional changes of ZFTA-RELA cells in monoculture versus coculture. **(i)** Cell state plots showing ZFTA-RELA tumor cells profiled by scRNA-seq and colored by culture condition. ZFTA-RELA patients profiled by sc/snRNA-seq were combined as a reference for cell state hierarchy. **(j)** Violin plots showing metaprogram scores for selected metaprograms in ZFTA-RELA cells cultured in monoculture or coculture. Boxplots denote Tukey’s whiskers (25–75 percentile represented by minima-maxima; statistical median as center). *P* values calculated by two-sided student t-test. **(k)** Comparison of somatokinesis (measured in µm per hour) of ZFTA-RELA cells in monoculture and co-culture (*n* = 206 cells in monoculture and *n* = 235 cells in co-culture, acquired from *n* = 3 independent biological replicates). Individual data points with mean±SE are shown. *P* values calculated by two-tailed Mann-Whitney test.

We next investigated whether cellular states with distinct morphologies also have distinct cellular behaviors. For this, we characterized subcellular and cellular tumor cell behavior with longitudinal live-cell imaging *in vivo* and *in vitro* co-culture models. Interestingly, neuronal-like cells were the most invasive state, followed by neuroepithelial-like cells, whereas EPN-glial-like cells were mostly stationary (**Figure 5c**). Moreover, we noted distinct modes of invasion across cell states. While neuroepithelial-like cells more frequently utilizing a continuous invasion mode, neuronal-like cells resembled immature neurons during neurodevelopment, more frequently adopting saltatory invasion patterns (**Figure 5d-e**, **Extended Data Figure 9a-**b)^43^. We therefore characterized the invasion patterns of neuronal-like cells in closer detail. We examined tumor TM dynamics, and observed behaviors reminiscent of neurites during neurodevelopment, including protrusion, retraction, and branching (**Figure 5f, Extended Data Figure 9c**)^44,45^. In addition, we distinguished three modes of neural-like cell migration, consisting of branching migration, locomotion, and translocation^25^ (**Figure 5g, Extended Data Figure 9d**). Overall, both on a subcellular and cellular level we observed how neural and neurodevelopmental invasion mechanisms are leveraged by neuronal-like ST-EPN cells to invade through their neural microenvironment of the brain.

Finally, we measured whether different cellular states have different proliferation rates. Neuroepithelial-like cells were the most proliferative compartment, which was consistent with our sc/snRNA-seq results (**Extended Data Figure 9e-f**), and with the expected behavior of immature progenitor cells. Neuronal-like cells, instead, were the least proliferative (**Extended Data Figure 9e**).

Taken together, these results suggest that ZFTA-RELA cellular states have distinct morphological and behavioral attributes in the context of their native neuroglial microenvironment, in which neuronal-like cell states are highly invasive neuroepithelial-like cell states are the most proliferative, and EPN-glial-like cell states are mostly stationary. These findings strengthen the concept of a division of labor across these cancer cell states, collectively cooperating to promote tumor progression.

### Neuroglial microenvironment induces tumor cell plasticity driving ZFTA-RELA tumor cell invasion

Having observed distinct morphology and behavior of tumor cell states in neural microenvironments, we asked whether brain-resident cells are necessary for inducing plasticity towards more mature cellular states. To do so, we compared our *in vitro* coculture models to monocultures consisting only of patient-derived ZFTA-RELA tumor cells (**Figure 5h**). We performed full length scRNA-seq of cells in coculture versus monoculture and found significant enrichment of neuronal-like GO terms and gene signatures only when tumor cells are grown in coculture, as well as a decrease in neuroepithelial-like signature (**Figure 5i-j, Extended Data Figure 10a-d**). These findings corroborate the morphological observations of less detection of EPN-glial-like and no detection of neuronal-like cells in monocultures (**Extended Data Figure 10e**). This suggests that normal brain-resident cells induce a shift of neuroepithelial-like progenitor cells towards neuronal-like lineages. Indeed, sc/snRNA-seq comparisons of three more ZFTA-RELA models grown as *in vitro* cultures (adherent and spheres) or in *in vivo* (patient-derived xenografts) revealed that indeed only ZFTA-RELA cells grown *in vivo* and cocultures are enriched in neuronal-like and ependymal-like states, demonstrating how brain-resident cells are required for plasticity towards these states (**Extended Data Figure 10f-g**). In accordance with our integrated molecular and functional cell state model, the overall invasion speed of ZFTA-RELA cells significantly increased due to a higher fraction of neuronal-like cell states in the coculture model (**Figure 5k**).

Overall, these results highlight how the neural microenvironment promotes plasticity towards a neuronal trajectory, affecting the morphology and behavior of ZFTA-RELA cells.

## Discussion

Our work uncovers different developmental signatures as well as malignant cell states across ST-EPN subgroups including previously uncharacterized non-canonical ST-ZFTA tumors^12^, and demonstrates that these cell states have distinct spatial, morphological, and behavioral characteristics.

Because of the limited number of ST-EPN patient samples available in previous studies^14,15^ and the recent re-classification of ST-EPN tumors into multiple molecular subgroups^12^, the etiology and cellular composition across ST-EPN tumors remained unknown. This ambiguity poses as a critical hinderance to creating precise and accurate stratification of patients, as well as effective personalized therapeutic decisions. Here, we introduce a comprehensive dataset of ST-EPN patient tumors and establish a compelling link between ST-EPNs and human embryonic cortical development. We highlight a clear discrepancy in both developmental signatures and differentiation lineages of different ST-EPN subgroups. While previous studies in mice have suggested radial glia as the candidate origin of ependymoma^12,46–50^, the human correlates of supratentorial ependymoma origins, especially of non-canonical ST-ZFTA subgroups, have remained elusive. In our work, we pinpoint to two distinct temporally restricted progenitor populations– neuroepithelial and embryonic cells - present during the first weeks of human embryonic development, as the most progenitor-like populations of ST-EPN tumors. Notably, while ZFTA-RELA and ZFTA-Cluster 3 tumors both harbor the same *ZFTA-RELA* fusions, they possess distinct progenitor signatures. Expanding on previous studies which observed that different *ZFTA* fusions are sufficient to drive tumorigenesis in neonatal mouse forebrain cells^12^, our observations allude to the possibility of the same fusion oncoprotein hijacking transcriptional programs of two different temporally restricted progenitors. This oncogenic event likely drives distinct downstream biology and clinical implications, also suggested by their disparate histological features^12,51^, gene signatures, and extent of differentiation to neuronal and/or ependymal differentiation. In contrast, ZFTA-Cluster 1 tumors harbor the *ZFTA-RELA* fusion and share a common neuroepithelial-like progenitor population with ZFTA-RELA tumors but lacks neuronal differentiation. This suggests differential maturation block for the two subgroups following fusion-driven transformation of the same progenitor. We therefore observe two separate oncogenic phenomena involving *ZFTA-RELA* fusion across subgroups, which would be of interest to validate further in patient-derived and mouse models. In addition, we validate previous findings that tumors harboring different fusions are distinct entities^8,12,13,52–54^. We highlight how ZFTA-Cluster 2, ZFTA-Cluster 4, and ST-YAP1 are transcriptionally and developmentally distinct from ZFTA-RELA tumors. In addition, the strong ependymal signatures in ST-YAP1 corroborates previous work demonstrating ciliogenic marker expression in these tumors^14,55^. These distinct developmental signatures and perturbed lineage differentiation across subgroups will have important therapeutic implication for developing differentiation therapy in ST-EPN cases.

Building on our description of molecular cellular states across ST-EPN tumors, we characterized how cellular states are spatially organized within ZFTA-RELA tumors. To our knowledge, no study has yet characterized spatial organization and the microenvironment of supratentorial ependymomas at single-cell resolution. To this end, we generated 10X Xenium data from 23 primary and recurrent ZFTA-RELA tumor sections, preserving morphological features of malignant cells. We first highlight hypoxia as a central driver of structural organization and demonstrate that highly vascularized tumor regions are more globally disorganized, implying that well-perfused tumor regions lack hypoxia-driven gradients of metabolic and phenotypic changes as observed in other systems^56–58^. Interestingly, this was recently described in adult gliomas by Greenwald et al.^24^, highlighting this as a potential common mechanism across brain tumors. Specific to ST-EPNs, we observe that relapsed patient tumors tend to have more MES/hypoxic regions and to be structurally more organized, pointing to hypoxia and structural organization as key attributes of a clinically aggressive subset of ZFTA-RELA tumors. Despite hypoxia being described as critical for metabolism and epigenetic regulation in posterior fossa ependymomas^59^, its role in ST-EPNs has been underappreciated. It would be of interest for future studies to examine hypoxia-driven tissue structural changes and its functional implications in ST-EPN tumors. In addition to global tumor spatial patterns, local tumor structures are predominantly enriched by the same cellular state, also described as “state-specific clustering” in gliomas^24^. This common phenomenon across brain tumors suggests spatial location as a key regulator of cellular state.

The neural microenvironment has been increasingly recognized for its role in driving tumor growth and invasion^17–20^. In addition, cell morphology and behavior are critical sources of cellular heterogeneity, and historically have been used to distinguish between cells across disciplines^60–62^. However, the integration of molecular cell states and their dynamic behavior in the native neural microenvironment have yet to be examined for ST-EPN tumors. As such, by integrating morphology and immunofluorescence with spatial transcriptomics and live cell imaging we demonstrate the molecular and phenotypic distinction between tumor cellular states. We find that neuronal-like ZFTA-RELA cells display long TMs and adopt highly invasive behavior, reminiscent of immature neurons during development. These neuronal-like cells with neuronal morphology and behavior have also been described in other types of gliomas^25^, highlighting a similar invasive tumor subpopulation in ST-EPNs tumors. This is notable, as ependymoma is thought to grow mostly locally at first and only show invasion at relapse, oftentimes increasing with the number of relapses^63^. This warrants investigation into tumor material taken at the border of resection in patients to further explore the role of this tumor subpopulation in tumor invasion. Furthermore, neuroepithelial-like cells undergo high number of cell divisions, being a candidate driver of proliferation within the tumor. These two distinct behaviors of cellular states suggest division of labor across malignant subpopulations, where neuronal-like cells drive invasion while neuroepithelial-like cells maintain proliferation. This functional specialization within tumors has been modeled in various studies^64,65^ and points to how transcriptional heterogeneity may lead to cellular cooperation in ST-EPN tumors.

Finally, while studies have described the synaptic connections between tumors and neurons^17,19^ as well as the effect of brain tumors on the neural microenvironment^66,67^, the role of the neural microenvironment on tumor cell plasticity has yet to be studied. In our work, we uncover how the neural microenvironment shapes the plasticity of ST-EPN tumor cell states towards more mature neuronal trajectories. We demonstrate that this plasticity leads to overall more invasive cell behavior, highlighting how the neural microenvironment not only influences cell plasticity but also subsequent tumor cell behavior. Our findings thus suggest that the effect of the neural microenvironment on tumor cells is an important area for future studies. Overall, our integrated analysis framework could serve as a blueprint to understand different functions of tumor cellular states and their plasticity across cancer entities.

In conclusion, we have demonstrated that integrating extensive transcriptomic, spatial, morphological and cellular behavioral characterization of patient tumors on the single cell level results in uncovering novel signatures of ST-EPN tumors related to the mechanisms of invasion, proliferation and plasticity. Furthermore, this study provides a methodological framework to perform such multidimensional profiling to uncover underlying biology across tumor types.

## Methods

### Experimental model and subject details

#### Human subjects

All deidentified samples used in this study were obtained after informed consent of patients and/or their legal representatives who did not receive compensation. The study was approved by the institutional review board in agreement with local institutional ethics guidelines DFCI 10-417 (Boston Children’s Hospital and Dana-Farber Cancer Institute), S-531/2020 (Heidelberg University), and EK Nr. 1244/2016 (Medical University of Vienna). Cohort characteristics are provided in **Table S1**.

#### ZFTA-RELA patient derived xenografts

EP1NS, BT165 and VBT242 cells (500,000 cells in 3 μL PBS per mouse) were injected stereotactically into the cortex of 6-week-old female NSG mice (NOD.Cg-*Prkdc*scid *Il2rg*tm1Wjl/SzJ, The Jackson Laboratory, RRID: IMSR JAX:005557), which were treated with 0.05 mg/kg buprenorphine and anesthetized with 2% to 3% isoflurane. The skull of the mouse was exposed through a small skin incision, and a small burr hole was made using a 25-gauge needle at the selected stereotactic coordinates: 1.0 mm X, 1 mm Y and -1.5 mm Z. After injection, mice were checked daily for signs of distress, including seizures, weight loss, or tremors. Tumor size was monitored monthly by small animal MRI and BLI starting 4 weeks after injection. Mice were euthanized as they developed neurologic symptoms, including head tilt, seizures, sudden weight loss, loss of balance, and/ or ataxia. All animal studies were performed according to Dana Farber Cancer Institute Institutional Animal Care and Use Committee (IACUC)–approved protocols (13-053).

#### ZFTA-RELA patient-derived cell lines

EP1NS, BT165 and VBT242 cell lines were generated as previously described^69,70^ and grown in “complete Neurobasal medium”, which consisted in Neurobasal™-A Medium (ThermoFisher Scientific, 10888022) supplemented with 1X of Antibiotic-Antimycotic 100X (ThermoFisher Scientific, 15240062), 1X GlutaMAX^TM^ (ThermoFisher Scientific, 35050061), 1 mg/mL Heparin solution (StemCell Technologies, 7980), 1X B-27 (Thermo Fisher Scientific, 12587010), 20 ng/ml recombinant Human FGF-basic (Shenandoah Biotech, 100-146) and 20 ng/ml recombinant Human EGF (Shenandoah Biotech, 100-26). For further passaging, floating cells were centrifuged at 300g for 5 min, dissociated with Accutase (Innovative Cell Technologies, Inc., AT104-500) for 5 min at 37 °C, and washed with PBS (Gibco, 10010023).

### *In vitro* co-culture

As described previously^71^, primary cultures of rat cortical neurons and astrocytes were prepared from E19 embryos. They were plated at a density of 90,000 cells/cm^2^ on 12mm diameter coverslips on 24-well plates that were previously coated with poly-L-lysine. The cells were maintained in neurobasal medium supplemented with B27 supplement (50x, 2% v/v) and L-glutamine (0.5 mM). After 7 days, Accutase dissociated, cells from patient-derived ZFTA-RELA cell lines were added and co-cultured (1000 cells/well). For timelapse live-cell imaging or immunohistochemistry, coverslips were used 4-14 days *in vitro* after seeding ZFTA-RELA cells. As a control, *in vitro* monocultures of ZFTA-RELA cells were established from patient-derived ZFTA-RELA cell lines as follows. 12mm diameter coverslips on 24-well plates were coated using poly-L-lysine and maintained in neurobasal medium supplemented with B27 supplement (50x, 2% v/v) and L-glutamine (0.5 mM). After 7 days, Accutase dissociated, ZFTA-RELA cells were added and cultured (1000-10000 cells/well). For timelapse live-cell imaging immunohistochemistry, coverslips were used 4-14 days *in vitro* after seeding ZFTA-RELA cells.

#### Fresh tumor processing

Live cells were isolated from immediately processed fresh tumor tissue acquired at the time of surgery. Fresh tumor tissue was mechanically dissociated and enzymatically digested using the Brain Tumor Dissociation Kit (Miltenyi Biotec, 130-095-942) and gentleMACS dissociator (Miltenyi Biotec, 130-096-427) for 30 min at 37 °C. Single-cell suspensions were filtered through a 70 µm strainer (Fisher Scientific, 03-421-228), centrifuged at 500*g* for 5 minutes and resuspended in a solution of 1% bovine ser albumin (BSA) (Biolegend, 644710) in phosphate-buffered saline (PBS) for fluorescence activated cell sorting (FACS).

#### Frozen tumor processing

Nuclei for snRNA-seq were extracted from snap frozen as well as optimal cutting temperature (OCT)-embedded tumor tissues. Snap frozen tumor tissue was cut on ice while OCT-embedded tissue was cut on a cryostat at -20 °C and washed with PBS. Frozen tumor tissue was suspended in Cell Lysis Buffer (CST) and dissociated with sterile surgical scissors for 5 minutes until homogenous. Dissociated tissue was filtered through a 70 µm strainer, 1XST was added to the nuclei solution and once more transferred through a 70 µm strainer. The nuclei solution was centrifuged at 500*g* for 5 min. Single-nuclei suspensions were re-suspended in PBS supplemented with 1% BSA (Smart-seq2), or 0.05% BSA (10X Genomics). All steps were performed at 4 °C.

#### Live cell sorting

A single cell suspension of 100 μL (kept in a FACS-tube in PBS/BSA at 4°C) was used for all sorting procedures as unstained control. For stained controls 0.75μL Calcein AM (Life technologies, C34851) and 0.5μL TO-PRO3 Iodide (Life technologies, T3605) were added to resuspended cells in 100μL PBS/1%BSA for 10 min at room temperature (RT). Single-cell sorting was performed on a SH700 sorter (Sony). Single viable tumor cells were selected by positive staining for Calcein AM as well as negative staining for TO-PRO3 and sorted into pre chilled (4 °C) 96 well plates containing TCL buffer (Qiagen, 1031576) + 1% β-mercaptoethanol (Thermo Fisher. 21985023). Sorted plates were centrifuged at 1000 x g for 1 minute at 4 °C and frozen on dry ice followed by transfer to a -80°C freezer for long term storage before whole transcriptome amplification, library preparation and sequencing.

#### Nuclei sorting

Dissociated frozen tumor tissue derived single-nuclei suspensions were resuspended in PBS 1% BSA and stained with 2.5mM Vybrant DyeCycle Ruby Stain (Life Technology, V10309). Nuclei sorting was performed on a SH700 SONY sorter. Intact nuclei were sorted into pre-chilled (4 °C) 96-well plates containing TCL buffer + 1% β-mercaptoethanol. Sorted plates were centrifuged at 1000 x g for 1 minute at 4 °C and frozen on dry ice followed by transfer to a -80°C freezer for long term storage before whole transcriptome amplification, library preparation and sequencing.

#### Sorting of cells grown in cocultures

DIV 08 coverslips of mono- and co-culture were incubated with Trypsin for 5 minutes. Next, 10% FBS was added, and cells were washed from coverslips. The samples were stained with a DAPI as a dead cell marker to identify the live cell population. The single cell suspension was sorted with a FACSAria Fusion 2 (BD Biosciences). Tumor cells were identified by their tdTomato expression.

#### Full-length single cell/nucleus RNA sequencing

Smart-seq2 whole transcriptome amplification, library preparation and sequencing of single cells/nuclei were performed following the modified Smart-seq2 protocol as previously described^14,72–74^. RNA was purified with RNAClean XP beads (Beckman Coulter, A66514). Next Oligo-dT primed reverse transcription (RT) was performed using Maxima H Minus reverse transcriptase (Life Technologies, EP0753) and a template-switching oligonucleotide (TSO; Qiagen) followed by PCR amplification (20 cycles for scRNA-seq and 22 cycles for snRNA-seq) using KAPA HiFi HotStart ReadyMix (KAPA Biosystems, 07958935001), and by AMPure XP bead (Beckman Coulter, A63882) purification. Libraries were generated using the Nextera XT Library Prep kit (Illumina, FC-131-1096). Libraries from 768 cells with unique barcodes were combined and sequenced using a NextSeq 500/550 High Output Kit v2.5 (Illumina, 20024906) on a NextSeq 500 sequencer (Illumina).

### Single-cell RNA-seq data processing

#### Quality filtering

We aligned raw sequencing reads to hg19 genome by hisat2 (v2.1.0) and quantified gene counts using RSEM (v1.3.0) as raw counts. For Smart-seq2 data frozen tumors, we excluded cells with genes < 1000 and an alignment rate of < 0.1, while for fresh tumors filtering threshold was < 2500 genes and alignment rate of <0.2. We also removed genes with TPM > 16 in <10 cells.

#### Normalization and scaling

For the remaining good-quality cells and genes, we computed the aggregate expression of each gene as Ea(*i*) = log_2_(average (TPM*i*,1…*n*) + 1) and defined relative expression as centered expression levels, Er*i*,*j* = E*i*,*j* − average(E*i*,1…*n*). In total, 7873 high quality cells were retained frozen tumors, and 1093 for fresh tumors for downstream analysis. On average, we detected 3660 unique genes per cell in frozen tumors, and 5815 unique genes per cell in fresh tumors.

#### Identification of malignant and non-malignant cells

To discriminate between malignant and non-malignant cells, we used: (1) Seurat clustering, (2) unbiased cell type annotation using the automated annotation package SingleR (v1.6.1) using the Human Primary Cell Atlas^75^ data as a reference, and (3) inference of copy-number variation using the InferCNV R package (v1.8.0). Cells that showed presence of copy-number alteration, similar clustering patterns, and that were not classified by SingleR as macrophages, T-cells, or endothelial cells were classified as malignant. The other non-malignant cells were classified to their respective cell type lineage based on SingleR prediction and/or expression of known marker genes.

### Data harmonization, Louvain clustering and identification of differentially expressed genes

Graph-based clustering with data integration was adapted to independently identify cellular clusters and gene signatures. Highly variable genes (HVGs) were selected using Seurat (v5.0.1), and their relative expression values were employed for principal component analysis (PCA). To discern sample-specific biological variations (e.g., tumor-specific genetic and epigenetic alterations) from cell subpopulation-specific variations and integrate multiple samples, a linear adjustment method (Harmony v1.0)^76^ was applied to the first 100 principal components (PCs) with default parameters, resulting in a corrected embedding. The first 20 Harmony-corrected dimensions were chosen for uniform manifold approximation and projection (UMAP) embedding.

#### Defining Nonnegative Matrix Factorization (NMF) metaprograms

We performed NMF for each sample separately to capture heterogeneity per sample, and negative values were set to zero. We ran NMF using K=6, and derived NMF programs for malignant cells for each sample using the top 10,000 over-dispersed genes using PAGODA2 (v0.1.4) and selected the top 50 genes from each NMF factor and scored all malignant cells with these NMF programs. We clustered NMF programs of ST-EPN samples by hierarchical clustering (distance metric: 1-Pearson correlation, linkage:Ward’s linkage) on the scores for each NMF program. This revealed eight highly correlated sets of programs within ST-EPN tumors.

To annotate malignant ST-EPN cells, we scored cells/nuclei for the eight MP signatures (*G_j_*). We did this by calculating for each cell *i* a score *SC_j_*(*i*), quantifying the average relative expression (*Er*) of the top 30 genes within each MP (*Gj*) and comparing the score to the average relative expression of a control gene (*G_j_^cont^*): SCj(i) = average[Er(G_j_,i)] – average[Er(G_j_^cont^,i)]. The control gene-set was defined by binning all genes into 30 bins of aggregate expression levels (*Ea*) and randomly selecting 100 genes from the same expression bin for each gene in the gene-set *Gj*.

#### Cell state plot

We separated ST-EPN cells into neuroepithelial-like/embryonic-like versus neuronal-like/ependymal-like as previously described. In brief, we first calculated the y-axis value as *y = max(SC_neuroepithelial-like_,SC_embryonic-like_) - max(SC_neuronal-like_,SC_ependymal-like_)*. For neuroepithelial-like/embryonic-like cells (*y > 0*), the x-axis value was defined as x = log_2_(|*SC_ependymal-like_ – SC_neuronal-like_* |+1) and for neuronal-like/ependymal-like cells (*y < 0*), the x axis was defined as log_2_(|*SC_embryonic-like_ – SC_neuroepithelial-like_* |).

#### GO terms enriched in ST-EPN metaprograms

We performed gene ontology analysis to identify biological terms that are over-represented in each metaprograms. To this purpose, we used the top 30 metaprogram-specific marker genes (see above) as input for *enrichGO* function from the *clusterProfiler*^77^.

#### GO terms enriched in coculture versus monoculture

We performed gene ontology analysis to identify which pathways are over-represented in ZFTA-RELA cells cultured with rat cortical neurons and astrocytes (coculture) versus ZFTA-RELA cells in monoculture. To this purpose, we first used Seurat’s *FindAllMarkers* function (MAST test, log_2_ fold change > 0.5, minimally detected in 25% cells) to identify genes that are differentially expressed between the two conditions. We then used the *enrichGO* function from the *clusterProfiler*^77^ package by selecting the top 300 marker genes enriched in coculture versus monoculture.

### Generation of expression scores

#### Correlation to reference datasets

We used Seurat’s *RunPrestoAll* function to derive the top differentially expressed genes (adjusted *P*-value < 0.05) of cell types within two reference datasets of the developing human cortex^29,30^, and within ST-EPN molecular subgroups or cellular states. We then scored all ST-EPN malignant cells with marker genes identified across the developing human cortex, and other way around, using the scoring function described above to annotate malignant ST-EPN cells with MP marker genes.

#### Cycling scores

We used Seurat’s *CellCycleScoring* function to assign cell cycle scores. This function relies on gene signatures that have been previously shown to characterize S and G2/M cell cycle phases. We defined high cycling cells as cells with S-scores or G2/M scores > 0, whereas low cycling cells as cells with S-scores < 0 and G2/M scores < 0.

#### Validation of developmental signatures across bulk RNA-seq ST-EPNs

To validate the preferential mapping of ST-YAP1 versus ZFTA-RELA tumors onto ependymal, respectively neuronal cells, we downloaded a microarray dataset^13^ available on the R2 platform (r2.aml.nl; R2 internal identifier: ps_avgpres_gse64415geo209_u133p2) containing normalized gene expression matrix (Z-score) data from *n* = 49 ZFTA-RELA and *n* = 11 ST-YAP1 patient tumor samples. After removing *n* = 10 ZFTA-RELA and *n* = 2 ST-YAP1 samples (as they were present in our discovery snRNA-seq cohort), we scored samples using Seurat’s *AddModuleScore* function taking the NMF-derived gene signatures described above as input.

### Spatial transcriptomics with 10X Xenium

#### Gene panel design

For *in-situ* 10X Xenium analysis, a custom gene panel of 100 genes was designed to include metaprogram marker genes identified across different types of high-grade gliomas, including: (1) ZFTA-RELA patient tumors analyzed in this study (*n* = 35 genes); (2) diffuse midline gliomas^34^ (*n* = 10 genes); (3) glioblastoma^37^ (*n* = 27 genes); (4) genes that mark the same metaprogram in a combination of tumor types (intersection, *n* = 10 genes); and (5) normal cell types (*n* = 18) (**Table S7**). For ZFTA-RELA tumors, those genes encompassed the following metaprograms: cycling (*n* = 4 genes), neuroepithelial-like (*n* = 8 genes), radial glia-like (*n* = 3 genes), NPC-like (*n* = 12 genes), ependymal-like (*n* = 8 genes), MES-like (*n* = 4 genes); and 3 probes targeting ZFTA-RELA fusion transcripts. In addition, we employed the human brain pre-designed panel containing probes for 272 genes encompassing additional non-malignant cell types (oligodendrocytes, immune cells, endothelial, astrocytes, neurons, microglia, and VLMCs).

#### Fusion probe design

For detection of fusion transcripts, custom probes were designed for three different fusion sites between the exons of ZFTA and RELA (**Table S7**). 60 base pairs surrounding fusion site was used to derive junction-specific probes.

#### Sample preparation

For fresh frozen samples, tissue sections were cut at a thickness of 10µm with a Leica CM3050S cryostat and collected onto Fisherbrand™ Superfrost™ Plus microscope slides, followed by fixation and permeabilization. For FFPE blocks, 5 µm tissue sections on a Xenium slide underwent deparaffinization and permeabilization. A mix of pre-designed and custom gene expression probe hybridization occurred at 50°C overnight. After multiple washing to remove un-hybridized probes, ligation of probes and annealing of rolling circle amplification (RCA) primer occurred at 37°C for 2 hours. Circularized probes were then enzymatically amplified at 30°C for 2 hours, after which background autofluorescence was quenched chemically. Slide was then loaded into the Xenium Analyzer following nuclei staining.

#### Xenium *in-situ* imaging

Image acquisition and sample handling was automated within Xenium Analyzer for two slides per run. 15 rounds of fluorescence probe hybridization, imaging, and probe removal occurred within the Analyzer. Fast area scan camera with ∼200 nm per-pixel resolution was used for image acquisition, and Z-stacks were taken with 0.75 μm step size across tissue thickness. All Z-stack of images were then processed and stitched, using DAPI image as reference. Each fluorescently labeled oligonucleotide bound to amplified barcodes was detected and registered in each cycle. Unique optical signature from fluorescence intensity over the 15 rounds was used to identify a target gene. For cell segmentation, neural network was used to detect each nucleus from DAPI images.

#### Preprocessing of data

The preprocessing of Xenium platform data was conducted using the Seurat pipeline. Initially, the per-transcript location data, cell x gene matrix, cell segmentation, and cell centroid information provided in the Xenium outputs were imported using the *LoadXenium* function. Subsequently, SCTransform was employed for normalization, followed by standard procedures for dimensionality reduction and clustering.

#### Metaprogram cell assignment

To assign the sequenced cells an identity, we first clustered cells using Louvain’s algorithm and found clusters for resolutions between 0.1 and 1 with an increment of 0.1. We used SeuratWrappers’s *RunPrestoAll* function to identify genes that were differentially expressed. We only considered genes detected in at least 10% of the cells within each cluster and showed a mean log difference of at least 0.25. We applied the Wilcoxon rank-sum test with Bonferroni correction for multiple testing and only kept genes with an adjusted P-value less than 0.05. For each resolution, the clusters’ scores were calculated individually and correlated with the scores of the programs defined in the base and custom panels. Finally, we manually assigned cell identities based on similarities. To confirm the correct cell assignment, we confirmed expression of malignant metaprogram and/or non-malignant cell type marker genes, as well as positive expression of ZFTA-RELA fusion-1 transcript in malignant compared to non-malignant cells.

#### Spatial coherence score

We defined the *spatial coherence score*, a measure of how structure or disorganized a tumor is, as follows. After annotating each cell with a malignant metaprogram or non-malignant cell type, the data was divided into 200 bins per row and column, totaling 40,000 spots per tumor section, each of which was analyzed to identify the cells it contained. To achieve this, a spot annotation program was developed based on the program with the highest proportion within that spot. To determine if a sample is organized or disorganized, a coherence score was calculated by examining each spot and its surrounding 3×3 square neighbors. If the neighbors were found to be like the spot annotation, a score of 1 was added; otherwise, a score of 0 was assigned. This process was carried out separately for each sample and metaprogram. The scores for each program were combined and divided by the number of spots associated with that program. The final coherence score for each sample was calculated by averaging all the scores calculated for each program across that sample and scaled within 0-1.

#### Spatial niche

Spatial niche analysis was performed for each tumor section in R using the *BuildNicheAssay*’s function from Seurat (v. 5) using the following parameters: k.neighbor = 20 and niches.k = 4. In short, this function considers the n = 20 spatially closest neighbors to each cell and count the occurrences of each cell type present in this neighborhood. It then uses k-means clustering to group cells that have similar neighborhoods together, resulting in n = 4 spatial niches for each tumor section. To identify niches that were consistently detected across multiple tumor sections, we first calculated the frequency of each cell type within each niche. In this case, myeloid, microglia and T-cells were grouped together under “immune”, and endothelial and VLMCs under “vascular”. We then clustered the results obtained for each niche by hierarchical clustering (distance metric: 1-Pearson correlation, linkage:Ward’s linkage). This revealed five highly correlated sets of niches that we termed “recurrent niches”.

#### Cell neighborhood

To assess whether cells from distinct programs exhibit a higher-than-expected proximity, we performed neighborhood enrichment analysis employing Squidpy (v1.3.1)^78^

### Experimental procedures

#### Hematoxylin & Eosin staining

Tissue slides were immersed in Hematoxylin for 20 mins and washed in distilled water (4x 1 min). After washing, slides were immersed in 70% and 95% ethanol for 3 mins, followed by Eosin staining for 2 mins. The slides were further dehydrated in serial ethanol incubation, including 95% ethanol (2x 30 secs) and 100% ethanol (2x 30 secs). Finally, slides were incubated in Xylene for 3 mins, and mounted with coverslip to dry for 30 mins before proceeding to imaging.

#### Immunofluorescence of ZFTA-RELA monoculture and coculture cells

Coverslips were fixed at DIVs 4-14 using 4% (w/v) paraformaldehyde (ThermoFisher, J61899.AP) for 10 minutes and washed with cold DPBS. Next, they were permeabilized using 0.2% Triton-X-100 (Sigma, 10789704001) in DPBS for 10 minutes and blocked with 10% (v/v) fetal bovine serum (FBS) for 10 minutes. Following, primary antibodies (Beta-III-tubulin, Abcam, ab7751, 1:100 dilution; S100B, SySy, 287006, 1:200; Nestin, Abcam, ab22035, 1:100; Vimentin, Abcam, ab29547, 1:100; DCX, Abcam, ab18723, 1:100; CCDC40, Abcam, ab121727-1001, 1:100; GFAP, Abcam, ab7260, 1:100; RFP, SySy, 390004 or 409009, both 1:100) were applied in a 10% FBS solution for 1 hour at room temperature, shaking. Coverslips were washed 3× 5min using DPBS. Appropriate secondary antibodies (Alexa Fluor® 647, 568, 546 or 488, 1:500 dilution) were incubated for 1 hour at room temperature, shaking. After 3× 5 min washing with DPBS, coverslips were mounted using a “SlowFade Gold” solution and DAPI (1:10000 dilution in DPBS). Images were acquired at a widefield fluorescence microscope using a 20x objective with a numerical aperture of 0.7 and a pixel size of 541,66nm (Leica DM6000) or a numerical aperture of 0.8 and a pixel size of 324,78nm (Zeiss Axioscan).

#### Timelapse live cell imaging

Timelapse imaging was performed using either a widefield fluorescence microscope (Nikon Ti-HCS) with a 10x objective (NA 0.3) and 1.5x zoom or a confocal microscope (Zeiss Celldiscoverer7 with LSM 980) with a 20x objective (NA 0.7) and 0.5 zoom. Coverslips were imaged every 15 minutes for 8-24 hours at 37 degrees Celsius with 5% CO_2_. Images were acquired with a pixel size of 486,86nm (Ti-HCS) or 794,44nm (CD7).

#### Cell division quantification

Cell division was quantified using live cell time-lapse imaging in cocultures of neurons and ependymoma cells over 15-24 hours. Cells were assigned their morphological cell state at the beginning of each time-lapse and cell division events were manually counted.

#### Somatokinesis measurement

Somatokinesis measurement was performed using ImageJ/Fiji.^79^ Images were registered using the HyperStackReg plugin in ImageJ/Fiji. Cell somata were outlined and the center point of the cell was determined using the centroid function. The distance between the center points of two time points was divided by the experimental observation time.

#### Data and code availability

All raw and processed scRNA-seq and 10X Xenium datasets will be made available at the Gene Expression Omnibus (GEO) under accession number GSE260452 upon paper acceptance or reviewers’ request. The codes used for data processing and to reproduce the main figures in the paper will be made available on Github (https://github.com/FilbinLab/Multidimensional_STEPN.git) upon paper acceptance or reviewers’ request. All data needed to evaluate the conclusions in the paper are present in the paper and/or the Supplementary Materials.

## Acknowledgements

Some illustrations were created with BioRender.com. This work was supported by generous funding from: a Distinguished Scientist Award from the Sontag Foundation (to M.G.F.), a Career Award for Medical Scientist from the Burroughs Wellcome Fund (to M.G.F.), the NIH Director’s New Innovator Award (DP2) (to M.G.F.), a U54 ROBIN grant (to M.G.F.), a SPORE grant (to M.G.F.), Matthew Parker & Kelsey Rioux-Parker (to M.G.F.), Links Fore Liam (to M.G.F.). and Ependymoma Research Foundation (to M.G.F.). S.G.D. was supported by the Swiss National Science Foundation (grant number: P500PB_217787). K.K.M was supported by CERN Robert Connor Dawes Scientific Fellowship from the National Brain Tumor Society. Further, the work was supported by the Deutsche Forschungsgemeinschaft (DFG, German Research Foundation), SFB 1389, UNITE Glioblastoma, project ID 404521405 and project number VE1373/2-1516 (addressed to V. Venkataramani). V.V. received financial support came from the Else Kröner-Fresenius-Stiftung (2020-EKEA.135), Heidelberg University and the Research Seed Capital (RiSC) from Ministry of Science, Research, and the Arts Baden Württemberg. “Medical-Scientific Fund of and the Mayor of Vienna” (project # 14015 to J.G.), the “Verein unser_kind”, the ““Forschungsgesellschaft für Cerebrale Tumore”, and “Initiative Krebsforschung” Anniversary fund of the Austrian National Bank (project #16152 to J.G.). We thank G.A.V. Cruzeiro for assistance with live cell imaging, and Lisa Mayr for consultation of druggable targets.

## Author contributions

D.J. and S.G.D. conceived the study, designed the experiments, interpreted results, curated the data and wrote the manuscript with input from all co-authors. K.K.M. and D.R.G. generated DNA methylation and fusion data. S.T and E.R generated and analyzed co-culture experiments with the help of C.L.C. C.A.B, S.N, R.H, and C.M.N generated and analyzed 10X Xenium data. A.N, B.E, C.M.N, A.C.B, S.C, J.S.R, O.A.H, and M.L.S generated scRNA-seq data, and L.J contributed to the analysis. D.L.G, K.B, S.M.P, M.K, J.G, L.B generated resources. T.N and S.A provided consultation. K.W.P., V.V. and M.G.F. conceived and supervised the study, interpreted results, and wrote the manuscript with input from all co-authors.

## Materials & Correspondence

The study did not generate new unique reagents. The data generated in this study are available within the article and in its Supplementary Data Files. Correspondence and requests for materials should be addressed to Mariella G. Filbin.

