## Extended Data Figures 1-10 for "Single-cell multidimensional profiling of tumor cell heterogeneity in supratentorial ependymomas"

### Extended Data Figure 1

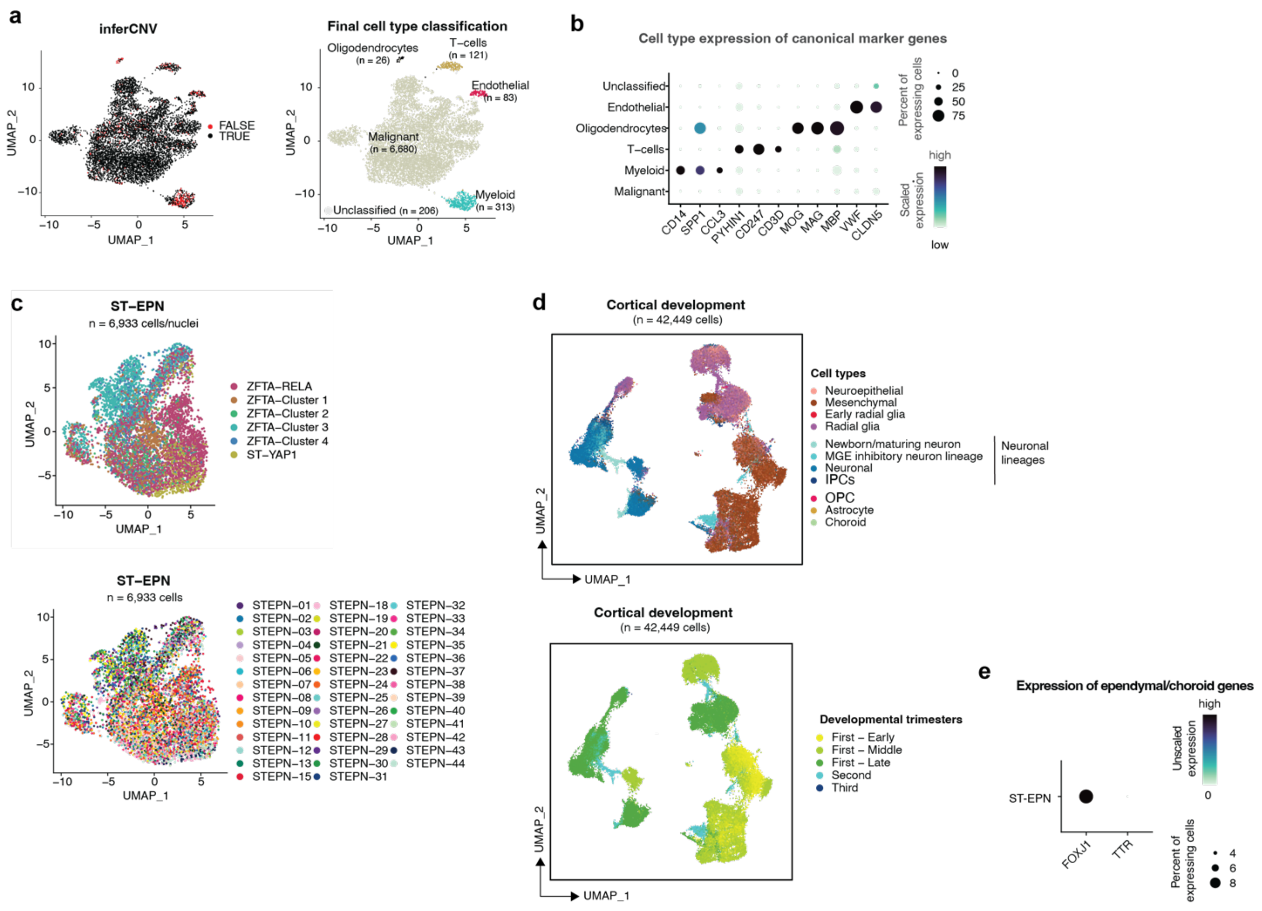


**Extended Data Figure 1. (a)** UMAP plot of all ST-EPN frozen patient nuclei (*n* = 7,429) profiled by snRNA-seq and integrated using Harmony[1]. Plots are colored by: inferred malignant status based on copy number variation (left panel) and final cell type classification (right panel). **(b)** Dotplot showing scaled mRNA expression of canonical marker genes of myeloid (*CD14, SPP1, CCL3*), T-cells (*PYHIN1, CD247, CD3D*), oligodendrocytes (*MOG, MAG, MBP*) and endothelial cell types (*VWF, CLDN5*) across all frozen ST-EPN cells profiled by snRNA-seq. **(c)** UMAP plot of malignant ST-EPN cells/nuclei integrated using Harmony[1]. Plots are colored by: ST-EPN molecular subtype (top panel) or sample of origin (bottom panel). **(d)** UMAP plot of cell types present during the first, second and third trimester of human cortical development[3,4], colored by cell type (top panel), and colored by developmental trimester (bottom panel) as specified in the original publication. First-Early: post-conception week (PCW) 4, First-Middle: PCW 5-6, First-Late: PCW 7-12. **(e)** Dotplot showing mRNA expression of *FOXJ1* and *TTR* across all frozen ST-EPN nuclei profiled by snRNA-seq. ST-EPNs show expression of the ependymal marker *FOXJ1* but not of the choroid marker *TTR*, suggesting ependymal projection.

### Extended Data Figure 2

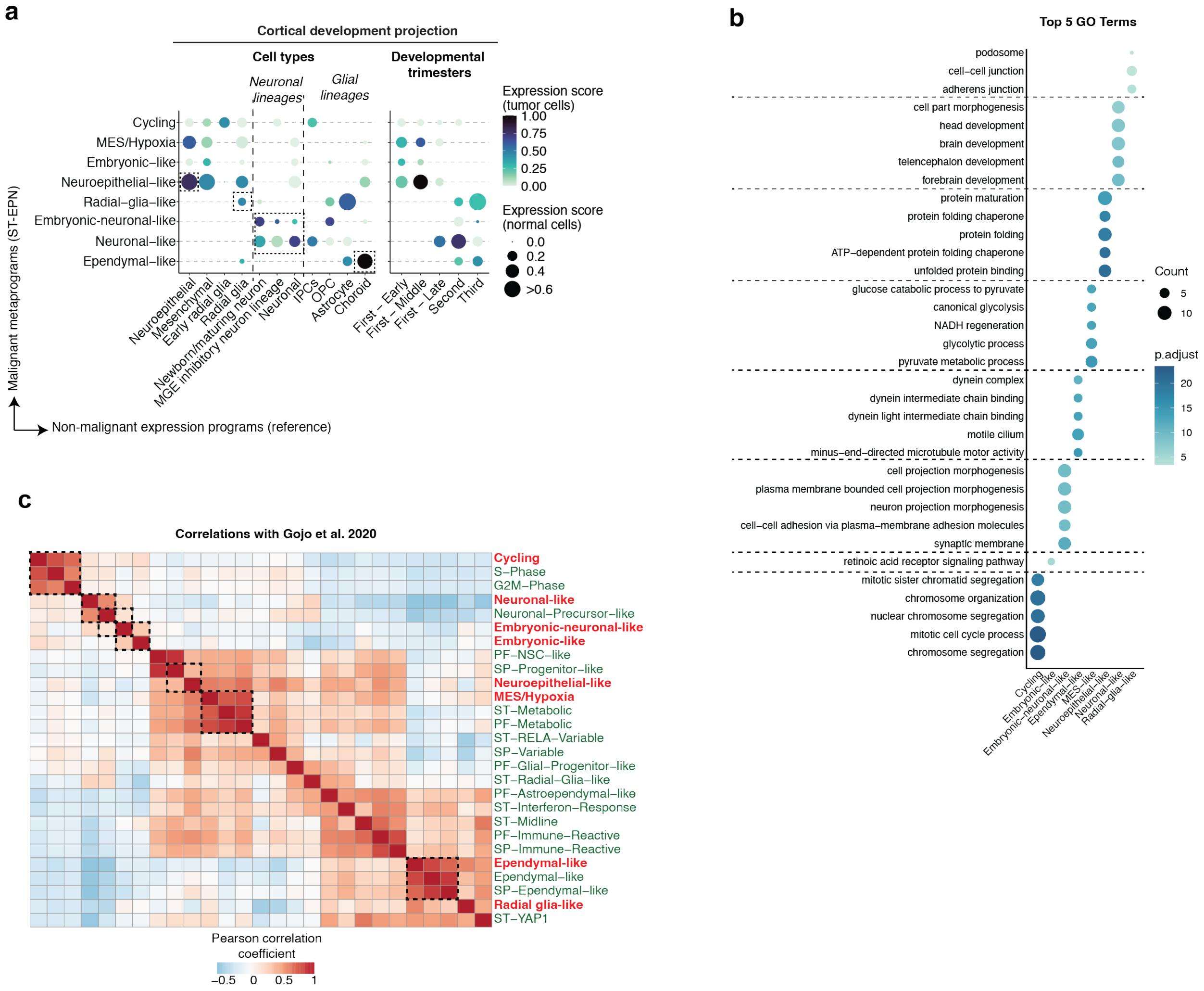


**Extended Data Figure 2. (a)** Projections of each ST-EPN metaprogram (y-axis) onto cell types present during the first, second and third trimester of human cortical development[3,4]. First-Early: post-conception week (PCW) 4, First-Middle: PCW 5-6, First-Late: PCW 7-12. Color scale depicts expression score of normal cell type gene expression in tumor cells, and dot size represents expression score of tumor cell type gene expression in normal cells. **(b)** Top five biological pathways enriched across the identified ST-EPN metaprograms by GO analysis. **(c)** Pairwise Pearson correlation between ST-EPN metaprograms identified in this study in frozen tumor samples (red font) and previously defined metaprograms[5] identified across different types of ependymoma (SP: spinal; ST: supratentorial; PF: posterior fossa).

**Extended Data Figure 3**

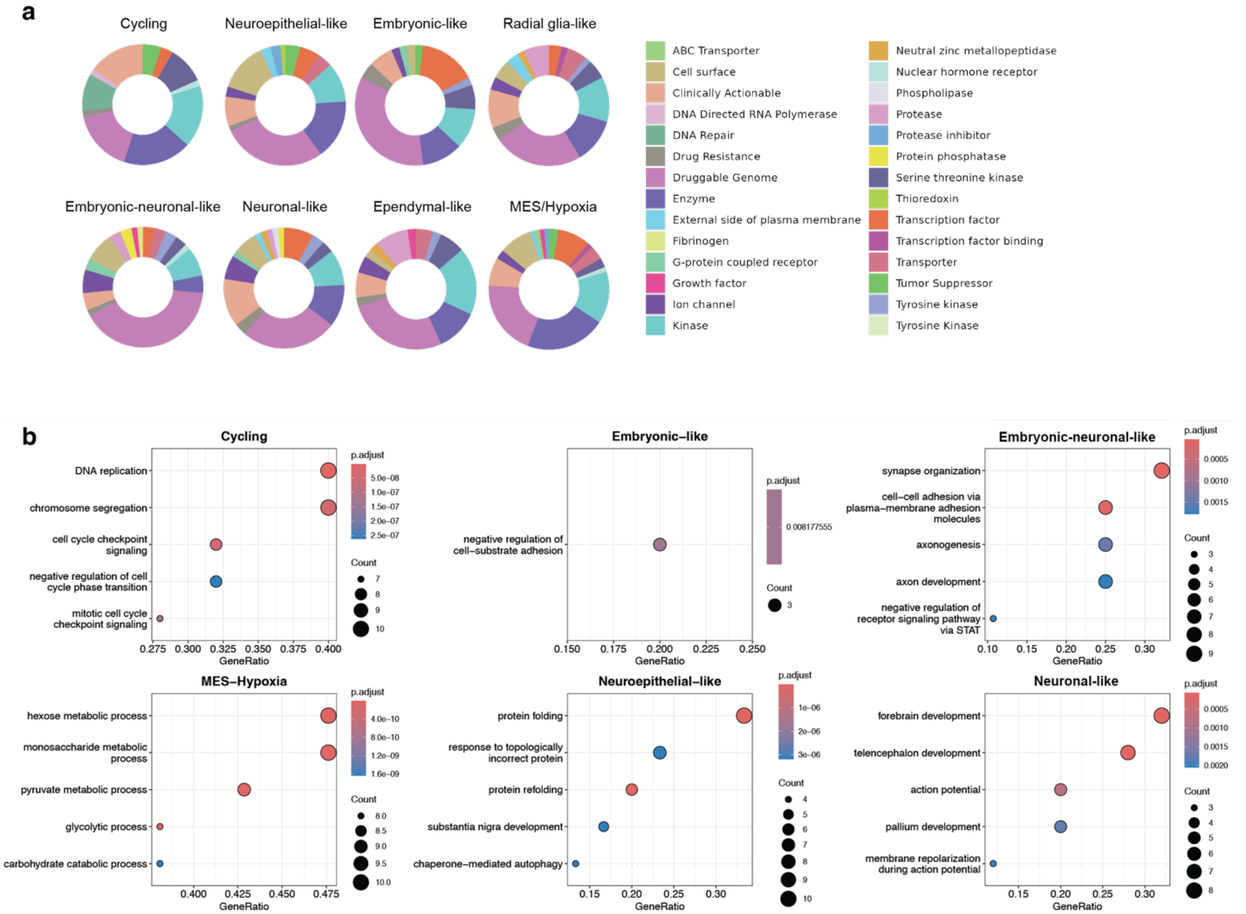


**Extended Figure 3. (a)** Pie charts of genes for therapeutic targets, analyzed by top 30 genes of ST-EPN tumor metaprogram integrated with DGIdb. **(b)** Top 5 GO terms of the ST-EPN tumor metaprogram gene hits in DGIdb ($p < 0.05$)

### Extended Data Figure 4

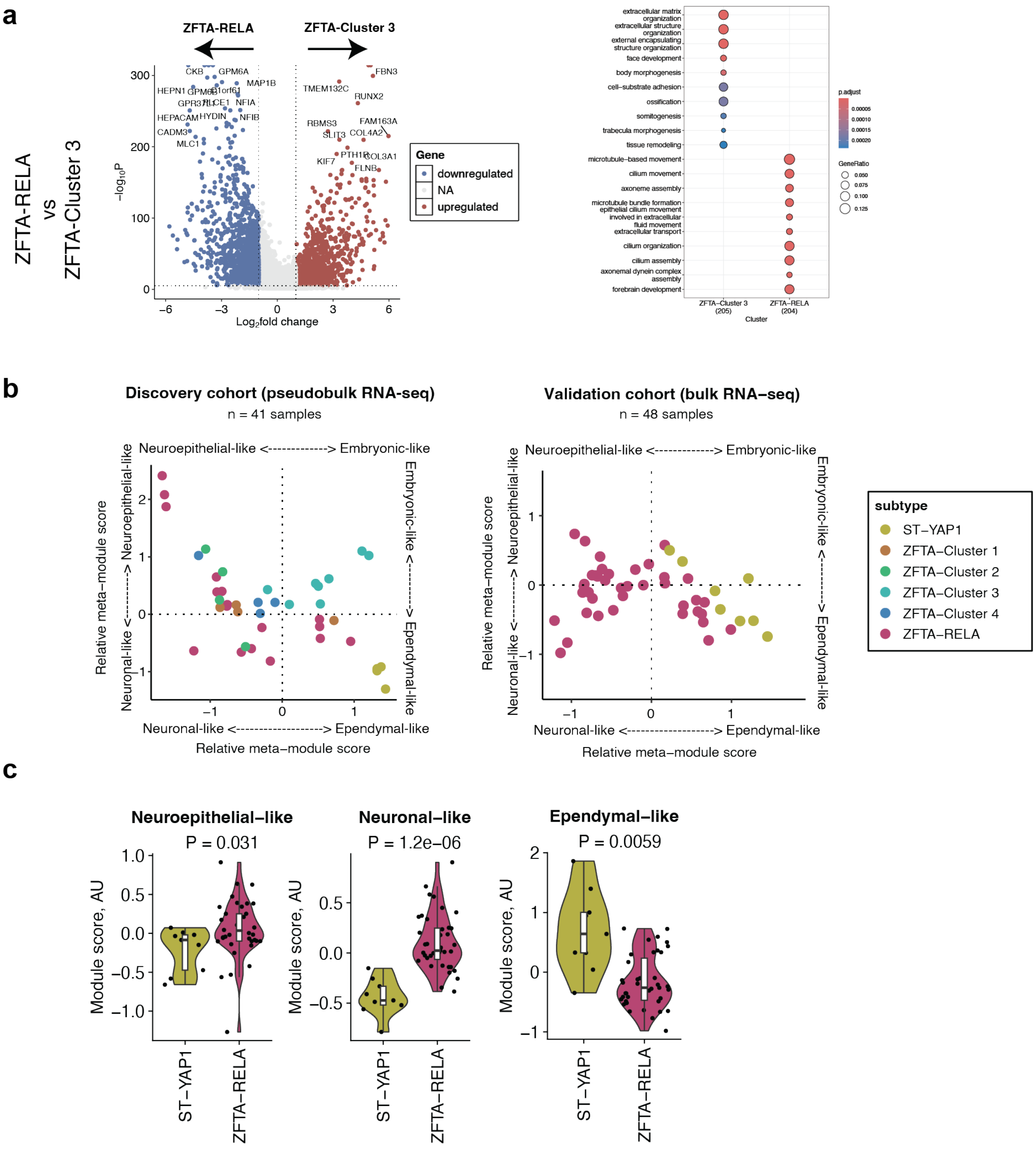


**Extended Data Figure 4.** Volcano plots depicting differentially expressed genes (left plot) and the top 10 enriched gene ontology terms (right plot) between ZFTA-RELA and ZFTA-Cluster 3 tumors. **(b)** Cell state plots of ST-EPN tumor cells/nuclei profiled in this publication and pseudo-bulked by sample of origin ("Discovery cohort", top) and of an external patient cohort[6] profiled by bulkRNA-seq ("Validation cohort", bottom). Samples were scored for metaprogram signatures identified by NMF analysis (top 30 genes) and colored by molecular subtype. **(c)** Violin plots showing gene expression module scores of the cell type signatures identified by NMF analysis across individual ST-EPN samples derived from the external cohort[13]. P-values calculated by student t-test are shown. AU, arbitrary unit.

**Extended Data Figure 5**

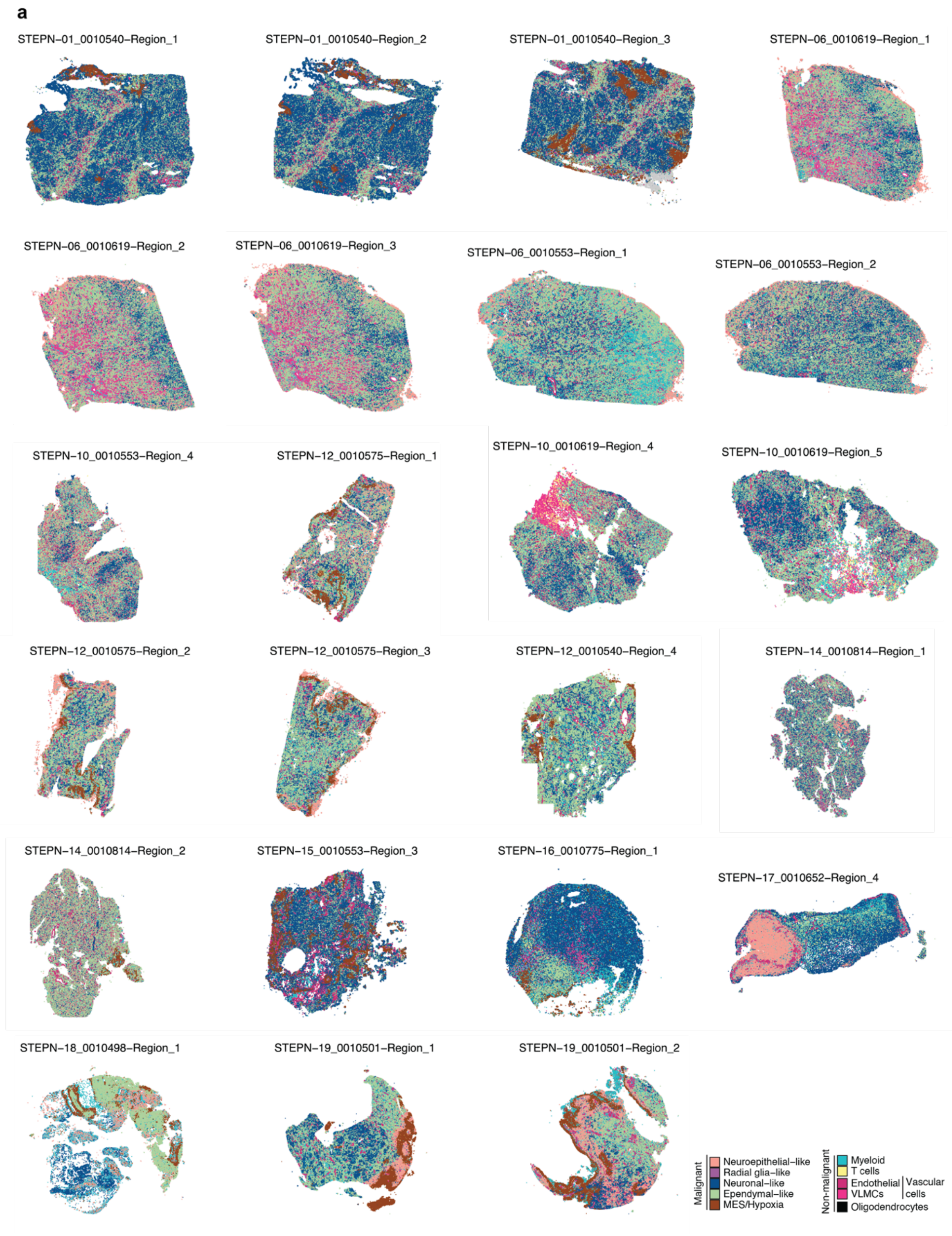


**Extended Figure 5. (a)** Spatial maps of 23 ZFTA-RELA tumor sections profiled by 10X Xenium, depicting distribution of both malignant metaprograms and non-malignant cell types.

### Extended Data Figure 6

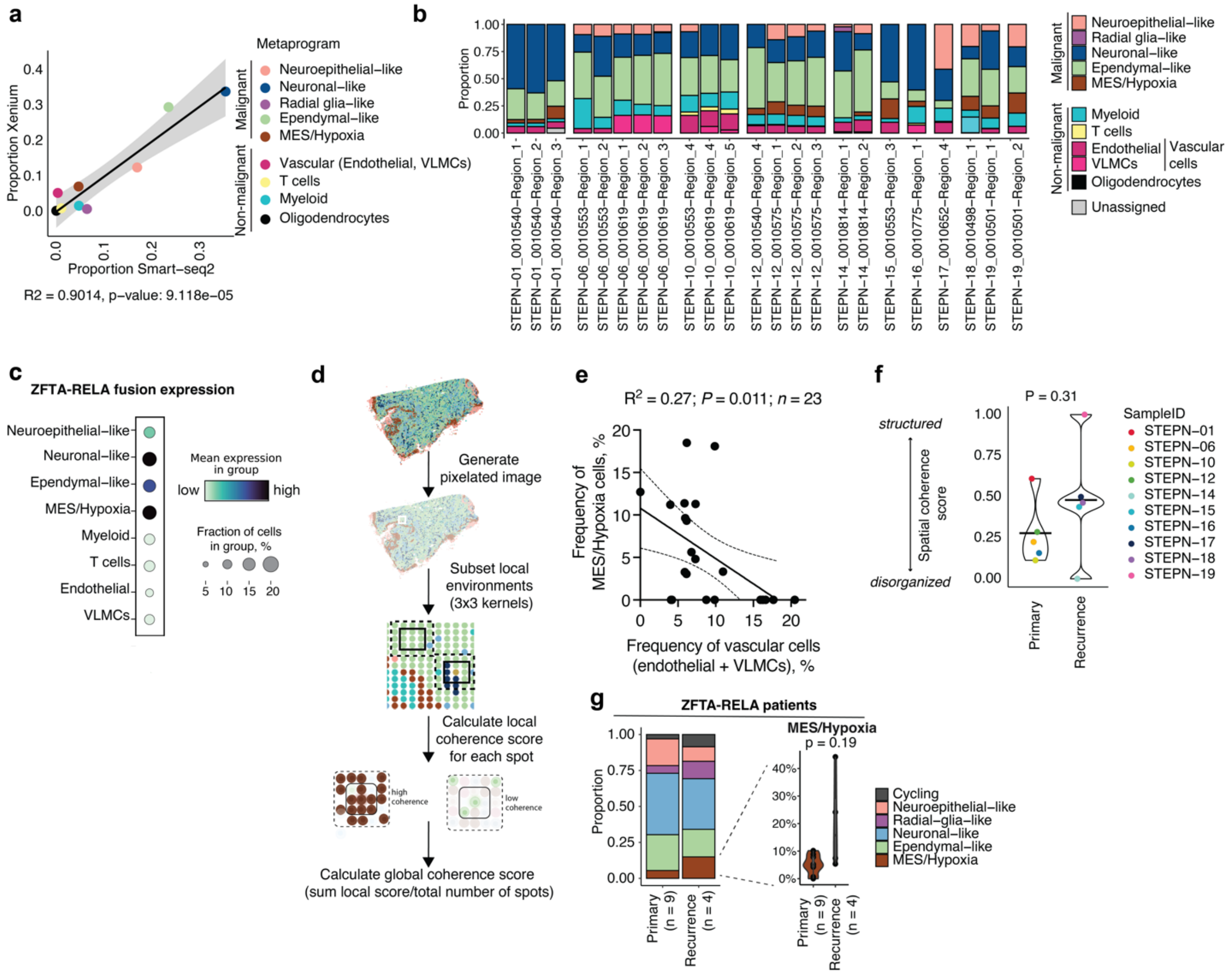

**Extended Data Figure 6. (a)** Scatterplot showing the frequency of cells detected in ZFTA-RELA tumors profiled by snRNA-seq and by 10X Xenium spatial transcriptomics. For the smart-seq2 dataset, neuronal-like and embryonic-neuronal like metaprograms were combined ("neuronal-like"), as well as neuroepithelial-like and embryonic-like ("neuroepithelial-like). The goodness of fit ($R^2$) and P-value are displayed at the bottom. **(b)** Proportion of tumor metaprograms and non-malignant cell types (y-axis) in all ZFTA-RELA tumor sections profiled by 10X Xenium (x-axis). **(c)** Expression of *ZFTA-RELA* fusion 1 transcript across the identified tumor metaprograms (malignant) and non-malignant cell types in all ZFTA-RELA patient samples profiled by 10X Xenium. **(d)** Schematics illustrating calculation of the spatial coherence score. **(e)** Correlation between the percentage of malignant MES/Hypoxia cells and non-malignant vascular cells. Data points are interpolated with a simple linear regression. Dashed line represents the 95% confidence intervals. The goodness of fit ($R^2$), P-value and number of data points are displayed at the top. **(f)** Spatial coherence across primary vs recurrent samples. *P*-value calculated by two-sided student *t*-test. **(g)** Mean proportion of ZFTA-RELA metaprograms across primary and recurrent samples profiled by sc/snRNA-seq. Neuronal-like and embryonic-neuronal like metaprograms were combined ("neuronal-like"), as well as neuroepithelial-like and embryonic-like programs ("neuroepithelial-like"). The percentage of MES/Hypoxia cells across individual samples is shown on the right. *P*-value calculated by two-sided student *t*-test.

### Extended Data Figure 7

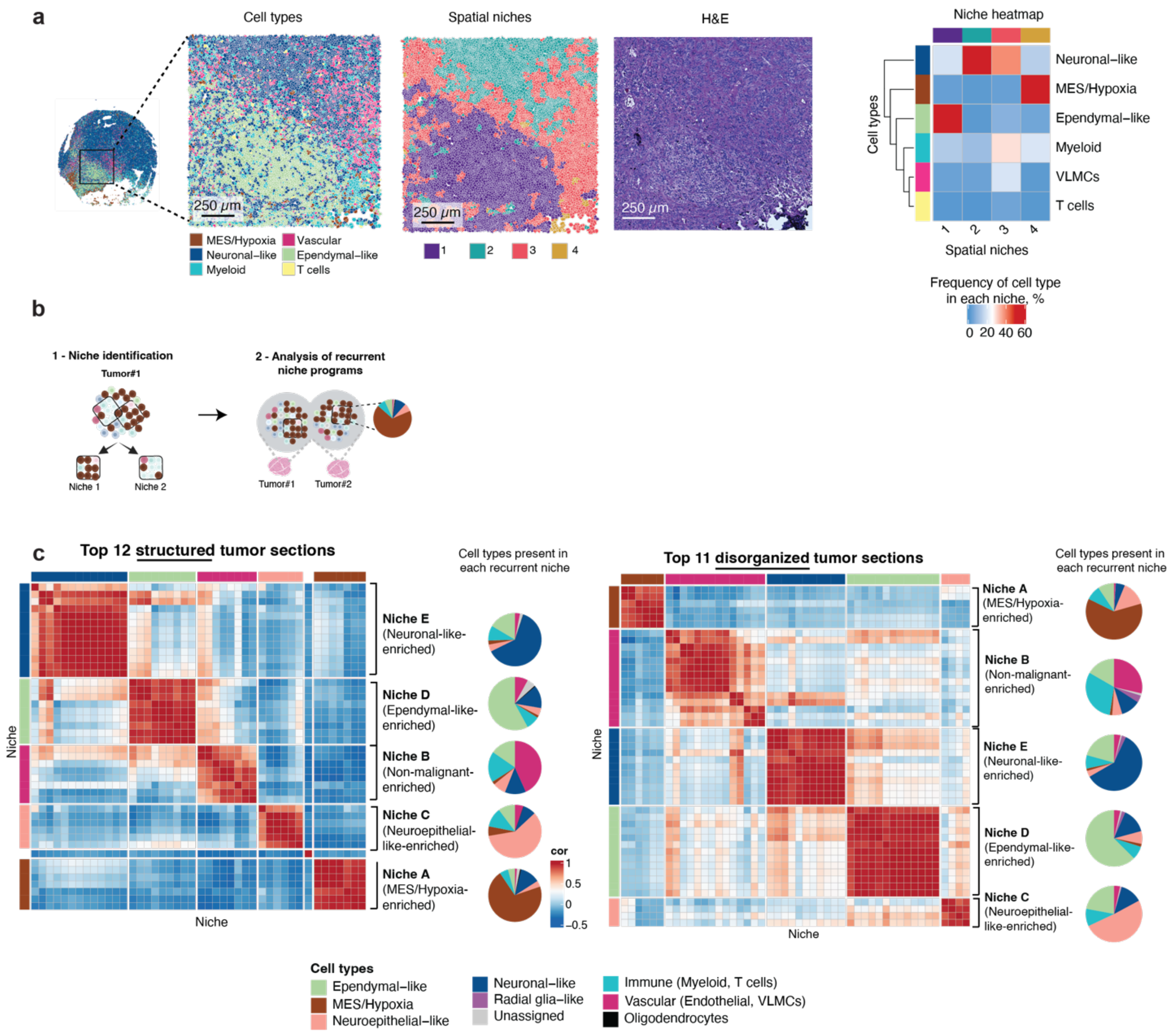


**Extended Data Figure 7. (a) (left)** Overlay of cell states/types, spatial niche, and H&E staining for a representative ZFTA-RELA tumor section (STEPN-16) showing morphological differences across regions and spatial niches corresponding to different tumor states. **(right)** Heatmap of corresponding sections depicting correlation between spatial niche (x-axis) and tumor metaprogram (y-axis), where color scale represents frequency of each cell type in each niche. (**b**) Schematic overview of the spatial niche analysis and identification of recurrent niche programs. (**c**) Recurrent niches identified split by tumor sections that are structured (left panel), and disorganized (right panel). Niches are annotated based on the most frequent cell type (displayed as pie charts on the right side of each heatmap).

### Extended Data Figure 8

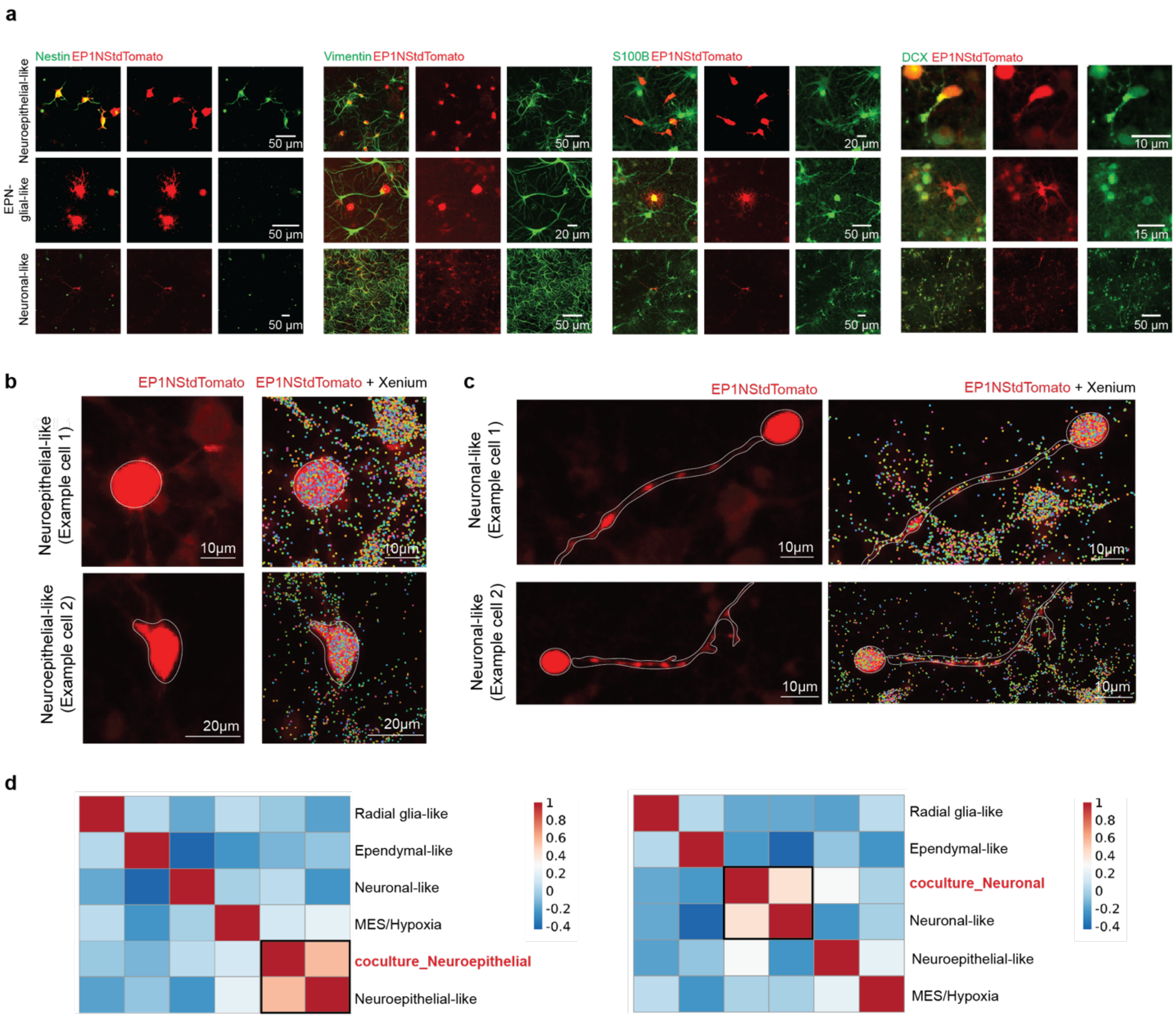


**Extended Data Figure 8. (a)** Representative immunofluorescence images of ZFTA-RELA (EP1NS cells expressing tdTomato) cells in co-culture, stained for different cell state markers: nestin and vimentin for neuroepithelial-like cell subtypes; S100B for EPN-glial-like cell subtypes; DCX for neuronal-like cell subtypes ($n$ = 234 cells acquired from $n$ = 3 independent biological replicates per marker). **(b and c)** Representative 10X Xenium analysis images of neuroepithelial-like **(b)** and neuronal-like **(c)** EP1NS-tdTomato cells in co-culture. Left panels depicts immunofluorescence images of tdTomato, and right panels represent Xenium transcripts overlayed with immunofluorescence image, where each dot represents detected transcript from the custom ($n$ = 100) and base panel ($n$ = 266) probes used for patient tumor sections. (**d**) Heatmap showing Pearson correlation between top 20 marker genes of tumor metaprograms and the top 20 marker genes of neuroepithelial-like cells ($n$ = 16 cells, left panel), and neuronal-like cells ($n$ = 16 cells, right panel).

### Extended Data Figure 9

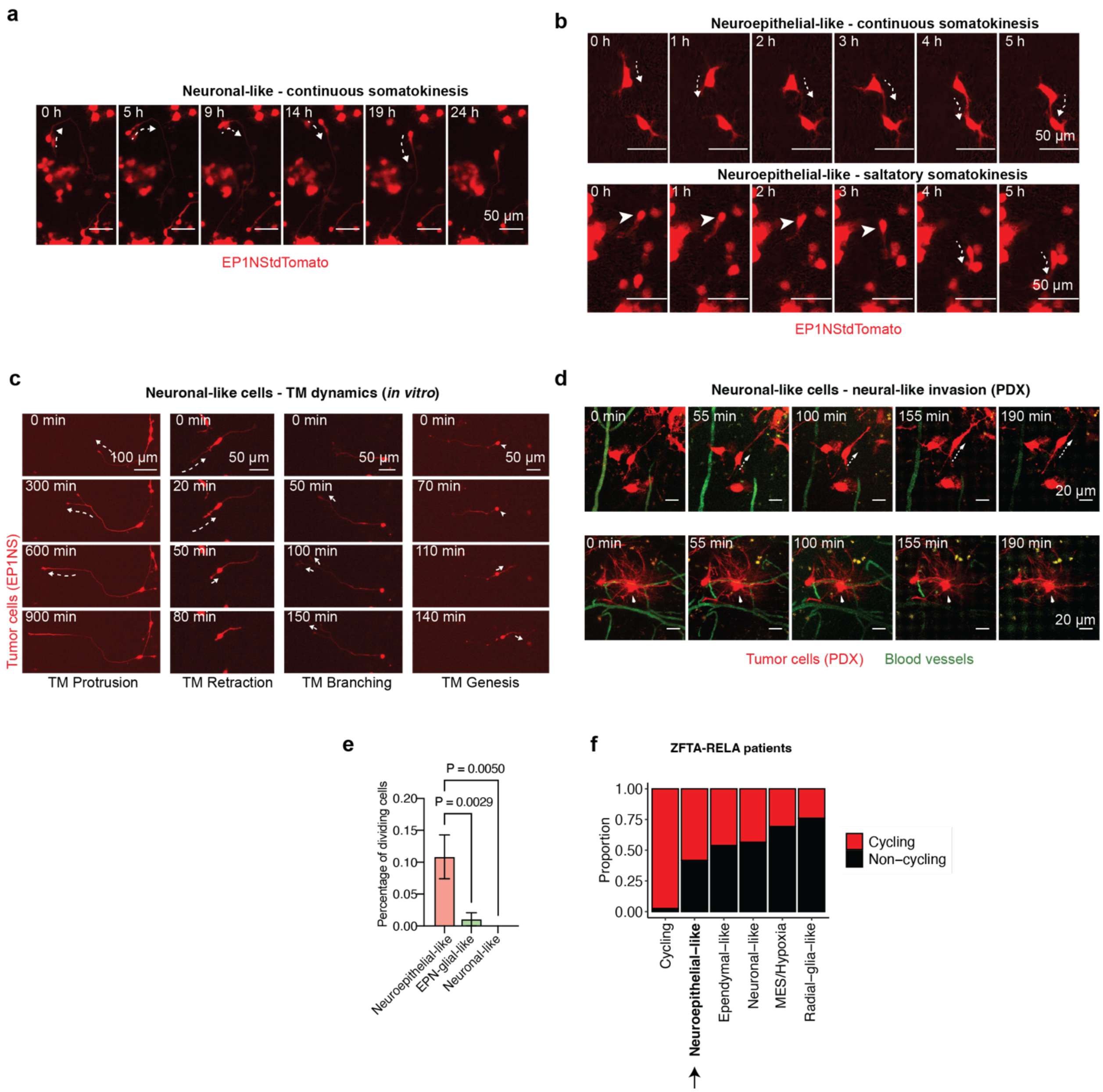


**Extended Data Figure 9. (a and b)** An exemplary invasive neuronal-like **(a)** and neuroepithelial-like **(b)** ZFTA-RELA tumor cell showing continuous migration in neural coculture. **(c)** Different forms of TM dynamics, such as protrusion, retraction, branching and TM genesis of an *in vitro* co-culture model of ZFTA-RELA (EP1NS cells) using live-cell timelapse imaging *in vitro*. Arrows indicate direction of TM movement. Arrowhead pointing to soma, out of which a new primary TM grows. **(d)** In-vivo time-lapse imaging of an *in vivo* PDX model of ZFTA-RELA (EP1NS cells expressing tdTomato) acquired with two-photon microscopy. Tumor cells are indicated in red, and blood vessels in green. An invasive cell and its invasion route (dashed arrow, above) versus a stable cell (arrowhead, below) are shown**. (e)** Percentage of dividing ZFTA-RELA tumor cells in coculture categorized by their morphological cell state (*n* = 83 neuroepithelial-like, n = 97 glia -like and n = 55 neuronal-like cells). one-way ANOVA with Tukey's multiple comparisons test. **(f)** Proportion of cells assigned as cycling or non-cycling (color-legend) by metaprograms in ZFTA-RELA patient samples profiled by sc/snRNA-seq.

### Extended Data Figure 10

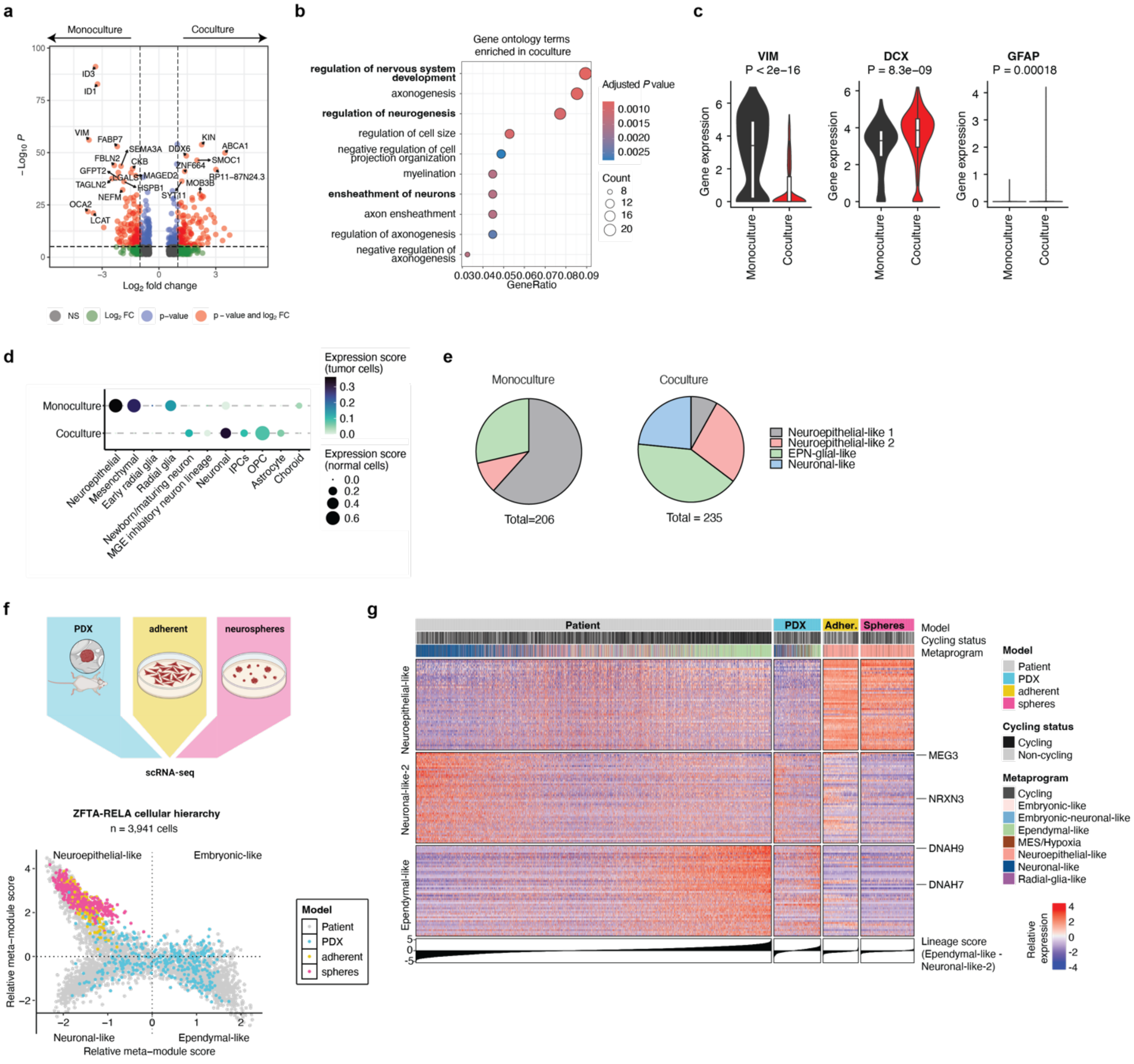

**Extended Data Figure 10. (a)** Volcano plot showing differential gene expression of ZFTA-RELA cells in coculture versus monoculture. Adjusted *P* values were calculated using MAST. Genes showing a $\log_2$ fold change > 1 and a *P* value < 0.00001 are highlighted in red. **(b)** Top 10 gene ontology terms of the marker genes upregulated in ZFTA-RELA cells in coculture versus monoculture. **(c)** Violin plots showing expression of *VIM* (neuroepithelial-like marker), *DCX* (neuronal-like marker) and *GFAP* (ependymal-like marker) in ZFTA-RELA tumor cells in monoculture and coculture profiled by scRNA-seq. Boxplots denote Tukey's whiskers (25–75 percentile represented by minima-maxima; statistical median as center). *P*-values calculated by two-sided student *t*-test are shown. **(d)** Projection of monoculture and coculture (y-axis) onto cell types present during the first, second and third trimester of human cortical development[3,4]. Color scale depicts expression score of normal cell type gene expression in tumor cells, and dot size represents expression score of tumor cell type gene expression in normal cells. **(e)** Morphological cell subtype distribution in ependymoma monoculture versus co-culture (*n* = 206 cells in monoculture, *n* = 235 cells in co-culture). **(f)** Graphical summary and cell state plot of ZFTA-RELA patient tumor cells profiled by sc/snRNA-seq overlayed with single cells derived from PDX (*n* = 3), adherent (*n* = 3) and neurosphere (*n* = 3) models of EP1NS, BT165 and VBT242. **(g)** Heatmap showing relative expression of the top 50 genes identified by NMF analysis specific to the neuroepithelial-like, neuronal-like-2 and ependymal-like signatures. Cells are sorted according to the overall expression of ependymal-like versus neuronal-like gene program. Model of origin, cycling status and metaprogram assignment are indicated on the top bars. Select genes of interest are indicated on the right.

**Extended Data Table legend.**

**Table S1.** Patient characteristics.

**Table S2**. Timepoint annotations used from Nowakowski et al. Science (2017) and Eze et al. Nature Neuroscience (2021).

**Table S3**. Cell annotations used from Nowakowski et al. Science (2017)

**Table S4.** Top 200 marker genes of NMF metaprograms identified in ST-EPN patients

**Table S5.** Gene ontology terms enriched in NMF metaprograms identified in ST-EPN patients (top 200 marker genes/metaprogram selected; GO terms with P-value < 0.01)

**Table S6**. DGIDB hits of NMF metaprograms identified in ST-EPN patients

**Table S7.** Genes included for spatial transcriptomics (Xenium)

**Table S8.** Proportion of metaprogram and cell types across ZFTA-RELA tumor sections analyzed by Xenium, and linear regression between spatial coherence score and proportion of selected metaprograms/cell types.
